# Suppression of NRF2-dependent cancer growth by a covalent allosteric molecular glue

**DOI:** 10.1101/2024.10.04.616592

**Authors:** Nilotpal Roy, Tine Wyseure, I-Chung Lo, Justine Metzger, Christie L. Eissler, Steffen M. Bernard, Ilah Bok, Aaron N. Snead, Albert Parker, Jason C. Green, Jordon Inloes, Sarah R. Jacinto, Brent Kuenzi, Benjamin D. Horning, Noah Ibrahim, Stephanie Grabow, Harit Panda, Dhaval P. Bhatt, Soma Saeidi, Paul Zolkind, Zoe Rush, Kathleen Negri, Heather N. Williams, Eric Walton, Martha K. Pastuszka, John J. Sigler, Eileen Tran, Kenneth Hee, Joseph McLaughlin, Géza Ambrus-Aikelin, Jonathan Pollock, Robert T. Abraham, Todd M. Kinsella, Gabriel M. Simon, Michael B. Major, David S. Weinstein, Matthew P. Patricelli

## Abstract

The NRF2 transcription factor is constitutively active in cancer where it functions to maintain oxidative homeostasis and reprogram cellular metabolism. NRF2-active tumors exhibit NRF2-dependency and resistance to chemo/radiotherapy. Here we characterize VVD-065, a first-in-class NRF2 inhibitor that acts via an unprecedented allosteric molecular glue mechanism. In the absence of stress or mutation, NRF2 is rapidly degraded by the KEAP1-CUL3 ubiquitin-ligase complex. VVD-065 specifically and covalently engages C151 on KEAP1, which in turn promotes KEAP1-CUL3 complex formation, leading to enhancement of NRF2 degradation. Previously reported C151-directed compounds decrease KEAP1-CUL3 interactions and stabilize NRF2, thus establishing KEAP1_C151 as a tunable regulator of the KEAP1-CUL3 complex and NRF2 stability. VVD-065 inhibited NRF2-dependent tumor growth and sensitized cancers to chemo/radiotherapy, supporting an open Phase I clinical trial (NCT05954312).

## Introduction

The NRF2 transcription factor (encoded by the *NFE2L2* gene) functions as a primary cellular defense against oxidative and electrophilic stress (*1, 2*). In the absence of cellular stress, NRF2 expression is restricted at the protein level by the Kelch-like ECH-associated protein 1 (KEAP1) and Cullin3 (CUL3) E3-ubiquitin ligase complex. KEAP1 functions as the substrate adaptor protein on the CUL3 scaffold, directly binding and positioning NRF2 for ubiquitylation. Hundreds of cellular stressors, including reactive oxygen species and electrophilic xenobiotics, lipids and metabolites, inactivate KEAP1-CUL3-mediated NRF2 degradation. Accumulated NRF2 protein then enters the nucleus to activate the transcription of ∼300 cytoprotective genes. Reactive cysteine residues in KEAP1 sense electrophilic stress, but precisely how this alters KEAP1 function to suppress NRF2 degradation is poorly understood (*3–5*).

Cancer evolution favors NRF2 activation where it promotes redox homeostasis, metabolic reprogramming, immune suppression, and resistance to chemotherapy, radiotherapy, and immune checkpoint inhibitors (*6–8*). Gain-of-function ‘hotspot’ mutations in NRF2 and loss-of-function mutations in KEAP1 or CUL3 are common in lung and upper aerodigestive cancers, resulting in constitutive NRF2 transcriptional activity (*1*). Mutation-independent mechanisms of NRF2 activation are also common, including KEAP1 post-translational modifications and altered protein-protein interactions that sterically displace NRF2 from KEAP1 (*1, 2, 9*). Characterization of these genetic aberrations and functional phenotyping of cells lacking NRF2 have provided compelling evidence in support of targeting NRF2 in cancer (*10–12*). Mouse models have also established that mutational activation of NRF2 functions in concert with hallmark oncogenes or tumor suppressors to enhance tumor initiation and progression (*11, 13, 14*). Conversely, genetic disruption of NRF2 suppresses tumor progression in transgenic mouse models of lung, pancreatic, and colorectal cancers (*9, 15*), strongly supporting NRF2 inhibition as a potential cancer therapy. Because NRF2 knockout animals are viable, fertile, and exhibit physiological disorders at older ages, NRF2 targeted therapies may be well tolerated (*16–19*). However, transcription factors such as NRF2 are challenging to drug, and to date, no direct NRF2 antagonistic therapies exist.

The KEAP1 protein includes 27 cysteines, several of which act as sensors to detect oxidative stress as well as various electrophilic metabolites and xenobiotics (*20*). Electrophilic attack of these cysteines inactivates KEAP1-mediated NRF2 degradation (*21, 22*). Cysteine-151 (C151) is the most extensively characterized sensor cysteine in KEAP1 and has been the subject of extensive drug discovery efforts (*23, 24*). FDA-approved KEAP1 inhibitors, such as dimethyl fumarate for multiple sclerosis and omaveloxolone for Friedreich’s ataxia, inhibit KEAP1 function through covalent modification of C151 (*25, 26*). In search of additional structurally distinct classes of KEAP1 inhibitors, we initiated a chemoproteomics screening-based drug discovery campaign aiming to identify KEAP1_C151 ligands that would activate NRF2 for treating auto-immune disorders. Surprisingly, however, during the optimization of our early leads, we discovered selective KEAP1_C151-reactive molecules that allosterically activate, rather than inhibit, KEAP1 via a novel molecular glue mechanism that promotes the KEAP1:CUL3 interaction, resulting in dramatically increased NRF2 degradation. Herein, we report the identification, optimization, mechanism of action and anti-cancer phenotypes for highly selective, potent and *in vivo* efficacious NRF2 inhibitors.

## Results

### Identification of covalent ligands that activate KEAP1

We utilized an industrialized targeted mass spectrometry (MS)-based chemoproteomics platform to screen a custom library of several thousand electrophilic small molecules for reactivity with several hundred cysteine residues, including KEAP1_C151 in native cells or cell lysates. This method measures changes in the covalent binding of cysteine residue to a biotinylated iodoacetamide probe following pre-treatment with candidate covalent ligands (*27, 28*). Molecules showing potent and selective KEAP1_C151 engagement in the chemoproteomics screen were assessed for functional impact on NRF2 using an engineered transcriptional reporter assay (HEK293-ARE-Luc), where luciferase is under the control of the antioxidant response element (ARE), a canonical binding site for NRF2. Two well-characterized KEAP1 inhibitors, PSTC and bardoxolone methyl, served as reference compounds which are known to increase NRF2 levels through covalent engagement to KEAP1_C151 (**Fig. 1A, S1A-C**). A simple morpholine acrylamide fragment, VVD-325, displayed sub-micromolar half maximal engagement potency (TE_50_) with KEAP1_C151 (**Fig. 1A & S1A**). Further chemical exploration of this chemotype led to VVD-330, showing a KEAP1_C151 TE_50_ equivalent to PSTC and bardoxolone, but, surprisingly, lacking cellular efficacy in the HEK293-ARE-Luc assay (**Fig. 1A & S1C**). To complement the HEK293 model, which has low basal NRF2 activity, we tested VVD-330 and its related analogues in KYSE70 cells, an esophageal squamous cell carcinoma (ESCC) cell line expressing a constitutively active NRF2^W24C^ mutation. Remarkably, VVD-330 reduced ARE-Luc activity in this NRF2 activated setting by 52% (**Fig. 1B**). The terminal phenyl group provided a useful handle for modulating the extent of NRF2 inhibition (I_max_), which proved to behave independently of engagement potency (**Fig. S1D**). For example, compounds VVD-446, VVD-860, and VVD-065 range from 45% to 94% I_max_ despite having very similar TE_50_ values (**Fig. 1A-B**). Removal of the acrylamide olefin in VVD-065 resulted in loss of activity in ARE-Luc assay (**Fig. S1E-F**), and time/dose response evaluation of VVD-065 engagement rates showed no signs of rate saturation at the highest dose tested of 25 μM supporting relatively weak reversible affinity (Ki > 25 uM) and a Kinact/Ki value of 3,497 M^-1^ s^-1^ (**Fig. S1G-H**).

**Figure 1:**
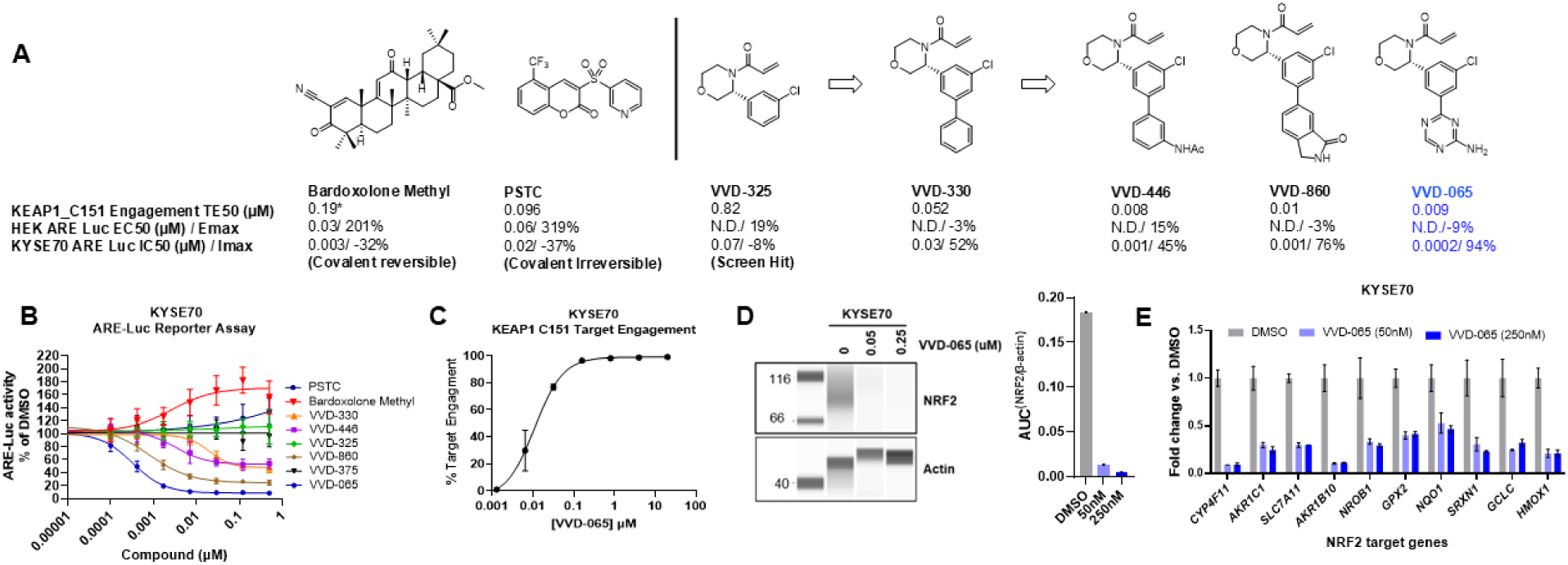
KEAP1 cysteine 151 liganding by VVD-065 induces NRF2 degradation and inhibits NRF2 target gene expression. (A) Progression of KEAP1_C151 ligands from the initial hit VVD-325 to more advanced VVD-860 and VVD-065. Bardoxolone methyl and PSTC served as reference compounds. (* Bardoxolone methyl is a covalent reversible compound and therefore the value is not directly comparable to the other values) (B) ARE-Luciferase reporter activity in NRF2^W24C^ KYSE70 cells treated for 18 hours with PSTC, Bardoxolone Methyl, and VVD KEAP1_C151 ligands (C) Chemoproteomic determination of KEAP1_C151 engagement by VVD-065 in KYSE70 lysates (D) Simple western analysis of NRF2 and Actin in KYSE70 cells treated for 18 hours with VVD-065 at indicated concentrations. Left: representative images Right: quantification of NRF2 normalized to Actin (E) Quantitative PCR based expression analysis of NRF2 target genes in KYSE70 cells treated for 18 hours with VVD-065 at indicated concentrations

VVD-065 emerged as an attractive lead compound from this campaign. It potently engaged KEAP1_C151 in KYSE70 cells (TE_50_ = 0.009µM; **Fig. 1C**), leading to robust degradation of NRF2 (**Fig. 1D**), inhibition of NRF2 transcriptional activity (**Fig. 1B**), and reduced expression of NRF2 target genes (**Fig. 1E**). Global chemoproteomic profiling confirmed exquisite selectivity of VVD-065 (**Fig. S1I-J; Table S1**), and its impact on NRF2 expression depended on both KEAP1_C151 (**Fig. S2A-B**), and proteasome activity (**Fig. S2C**).

### Structural and mechanistic evaluation of VVD-065 activity

To gain mechanistic insight into how VVD-065 affects KEAP1 function, we determined the crystal structure of the KEAP1 BTB domain in complex with this compound at 1.87 Å resolution (**Fig. 2A & S2D-E, Table S2**). As expected, the terminal carbon of the acrylamide of VVD-065 covalently binds C151. A hydrogen bond is formed between the side chain of R135 and the carbonyl of the acrylamide, and the morpholine ring projects into solvent, acting as a linker to spatially orient the reactive acrylamide relative to the binding cleft occupied by the biaryl group. The chlorophenyl moiety sits in a hydrophobic cleft and the triazine forms π-stacking interactions with H129 and H154. The terminal amine forms a water-mediated hydrogen bond with the backbone amide of H129 (**Fig. 2A**). Given the distance between the VVD-065 binding site and the KEAP1:NRF2 interface, it was not immediately clear how VVD-065 promotes NRF2 degradation (**Fig. 2B & S2D**). It has been suggested that binding of KEAP1_C151 inhibitory ligands in this groove decreases KEAP1:CUL3 interaction through steric displacement of the N-terminal peptide of CUL3 (*29*); however, this explanation is implausible because occupancy of the same binding site by VVD-065 promotes KEAP1 activity. Comparison of the KEAP1 BTB domain structure in the VVD-065 bound to unliganded or bardoxolone-bound state revealed major conformational changes induced by VVD-065 binding. The side chain of C151 is in the p rotamer when bound to VVD-065, contrasting to other published structures of both apo (**Fig. S2F**) and bardoxolone-bound (**Fig. 2C**) KEAP1 BTB domain where this residue is observed in the m rotamer. To accommodate the rotation of C151, M147 (beta-strand 5) and F52 (beta-strand 1) are pushed down and away from the BTB core. These movements change the hydrophobic packing of the BTB domain and enable helix 6 to pull in toward C151. The end of helix 6, defined by the α-carbon of Q177 in the truncated crystallization construct, moves by 2.9 Å. Importantly, the conformation of the KEAP1 BTB domain bound to VVD-065 can be nearly superimposed with the structure of the KEAP1 BTB domain when complexed with CUL3 (**Fig. 2D**). Both the shifted position of helix 6 and the p-rotamer of C151 are consistent between these two structures. The similarity between the KEAP1/VVD-065 and KEAP1/CUL3 complexes suggests that VVD-065 may increase KEAP1 activity by stabilizing a KEAP1 conformation that favors CUL3 binding.

**Figure 2:**
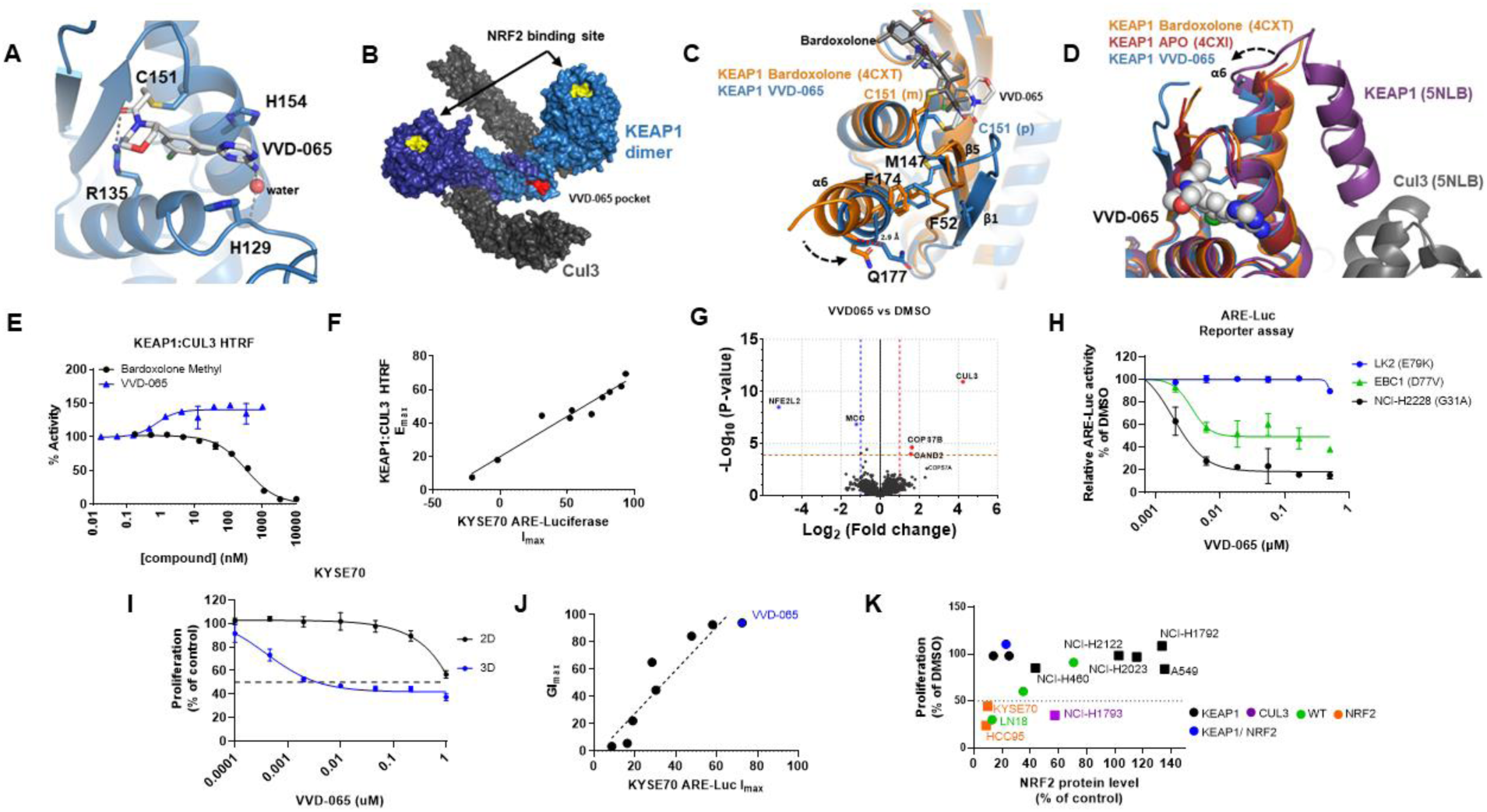
VVD-065 stabilizes KEAP1-CUL3 complex formation. (A) Binding interactions of VVD-065. VVD-065 binds in a hydrophobic cleft and forms a hydrogen bond with the sidechain of R135 and a water mediated interaction with the backbone amide of His129 (B) Model of the KEAP1-Cul3 complex highlighting the distance between KEAP1-NRF2 interface (yellow), the KEAP1-Cul3 interface and the VVD-065 binding site (red). KEAP1 (light and dark blue) forms a dimer through the BTB domain, and each copy independently binds to Cul3 (grey). VVD-065 binding site in KEAP1 monomer is shown in the figure (C) Superposition of bardoxolone bound KEAP1 (orange, 4CXT) with VVD-065 bound KEAP1 (blue) highlighting the concerted movement of residues C151, M147, F174 and F52 (D) Superposition of APO KEAP1 (dark red, 4CXI), KEAP1-Cul3 complex (purple and grey, 5NLB), KEAP1-bardoxolone (orange, 4CXT) and KEAP1-VVD-065 (blue). (E) Effect of Bardoxolone Methyl and VVD-065 on KEAP1 CUL3 interaction measured by HTRF assay (F) Correlation analysis between activities in KEAP1-CUL3 HTRF assay and KYSE70 ARE-Luciferase assay. E_max_= maximum increase in KEAP1-CUL3 interaction observed in HTRF assay; I_max_ = maximum inhibition of ARE-Luciferase activity in KYSE70 cells (G) MS analysis of KEAP1 interacting proteins in the presence and absence of VVD-065 using a miniTurbo based proximity labeling assay (H) ARE-Luciferase reporter activity in LK2 (NRF2^E79K^), EBC1 (NRF2^D77V^), and NCI-H2228 (NRF2^G31A^) cells treated for 18 hours with VVD-065 (I) Viability levels of KYSE70 cells treated with VVD-065 in either adherent (2D monolayer assay) or non-adherent (3D sphere assay) culture condition (J) Correlation analysis between activity in KYSE70 ARE-Luciferase assay (I_max_= maximum inhibition in ARE-Luciferase assay) and maximum growth inhibition (GI_max_)observed in 3D sphere assay (K) Correlation analysis of NRF2 degradation & growth inhibition in cells treated with VVD-065. NRF2 expression was measured using Simple Western. Growth inhibition at a concentration of 0.2 µM was plotted for all cell lines, except for NCI-H23 and KYSE180, where a concentration of 0.3 µM was used. Mutation status of each cell line is indicated by different colors. Cell lines showing >50% growth inhibition with genetic depletion of NRF2 (Fig. S3H) are indicated by filled square. Cell lines with <50% growth inhibition in Fig. S3H are indicated by filled circles

To test this hypothesis, we used HTRF (Homogeneous Time Resolved Fluorescence) assays to quantify protein-protein interactions (PPI) between recombinant KEAP1 and either CUL3 or NRF2. The KEAP1 activator VVD-065 increased KEAP1:CUL3 binding, while the inhibitor Bardoxolone decreased this interaction; neither compound affected KEAP1-NRF2 interactions (**Fig. 2E & S3A**). Evaluation of KEAP1 pre-engaged with VVD-065 in a competitive KEAP1-CUL3 HTRF assay confirmed the VVD-065-mediated increase in KEAP1-CUL3 affinity (**Fig. S3B**). The maximal impact on the KEAP1:CUL3 interaction (E_max_) correlated precisely with the maximal inhibition observed in the KYSE70-ARE-Luc assay, supporting that VVD-065 and its analogs promote KEAP1-mediated NRF2 degradation through enhanced KEAP1:CUL3 interactions (**Fig. 2F**).

To test whether KEAP1 activators like VVD-065 also promote KEAP1-CUL3 complex formation in live cells, KEAP1 proximal proteins were evaluated in the presence and absence of ligand using miniTurbo-based biotin proximity labeling. VVD-065 treatment strongly increased CUL3 within the KEAP1 biotinylation sphere (**Fig. S3C**). NRF2 was detected as a KEAP1-proximal protein in DMSO treated cells, but not VVD-065 treated cells, likely reflecting the overall loss of NRF2 in VVD-065 treated cell lysates. The impact was specific to CUL3 as the KEAP1 associated proteins P62/SQSTM1, MCM3, PGAM5, TSC22D4, and SLK (*30*) were unaffected by VVD-065 treatment (**Fig. S3C**). MS analysis further revealed strong increases in KEAP1-proximal CUL3, and additional proteins known to associate with the Cullin/RBX1 E3 ligase systems such as CAND2 (*31*) in response to VVD-065 treatment (**Fig.2G**).

Overall, these structural and biochemical studies. And cellular NRF2 degradation data indicate that VVD-065 binding to KEAP1_C151 leads to allosteric changes that increase KEAP1-CUL3 affinity and the abundance of ubiquitination-competent KEAP1-CUL3 complexes in cells.

### VVD-065 activity requires KEAP1-NRF2 interaction

Cancer-derived mutations in NRF2 localize to either the ^29^DLG or ^79^ETGE motifs, which are required for binding each monomer of the KEAP1 homodimer. The ^79^ETGE motif binds KEAP1 with ∼100-fold greater affinity than the ^29^DLG motif, supporting a model wherein ^29^DLG-mutant NRF2 retains KEAP1 binding via the ^79^ETGE, while mutations in the high-affinity ^79^ETGE motif result in a more profound loss of KEAP1 binding (*32*). As a tumor suppressor, KEAP1 mutations are not localized in hotspots and their impact on NRF2 binding is variable and complex (*33*). We explored the impact of VVD-065 on a set of cell lines with NRF2 mutations that either preserve or eliminate KEAP1-NRF2 binding: a) NRF2^W24C^ (KYSE70) and NRF2^G31A^ (NCI-H2228) that maintain KEAP1 interaction, b) NRF2^D77V^ (EBC1) that moderately compromises the association with KEAP1, and c) NRF2^E79K^ (LK2) in the ^79^ETGE motif that dramatically reduces binding to KEAP1 (*32*). Similar to what we observed in KYSE70 cells (**Fig. 1B**), VVD-065 induced a dramatic reduction of NRF2 transcriptional activity in NCI-H2228 cells (**Fig. 2H**). EBC1 cells showed a partial decrease in NRF2 transcriptional activity, while no impact was observed in LK2 cells, consistent with the relative KEAP1-NRF2 affinity of the two mutants expressed in these cell models. As expected, compound activity was seen in WT cell lines such as NCI-H520 and PC9 (**Fig. S3D**). Similar to NRF2^E79K^, the KEAP1^G333C^ mutation found in A549 cells completely disrupts KEAP1-NRF2 interaction (*33*), and VVD-065 failed to induce NRF2 degradation in these cells. CRISPR editing to convert the mutant *KEAP1* allele to WT restored VVD-065 suppression of NRF2 (**Fig. S3E**). We also confirmed that VVD-065’s functionality is independent of basal NRF2 activity level, as assessed by a NRF2 gene signature (**Fig. S3F-G**)(*7*). In summary, and consistent with our proposed mechanism of action, VVD-065 mediated NRF2 reduction requires the presence of KEAP1 and a residual ability of KEAP1 to interact with NRF2. These criteria are met in WT as well as a subset of KEAP1 and NRF2 mutant settings. It is to be noted that only such “mechanistic sensitive” settings, where VVD-065 degrades NRF2, are relevant for exploring the pharmacological effects of VVD-065.

### VVD-065 exhibits cancer cell growth inhibitory effects

Genetic inactivation of NRF2 impedes the growth of NRF2-active human cancer cell lines, and this NRF2 dependency is more profound under non-adherent culture conditions (*10, 12*). Consistently, VVD-065 inhibited the growth of KYSE70 cells in non-adherent, but not in adherent culture conditions (**Fig. 2I**). Across VVD-065 and a set of related analogs, we observed a strong correlation between ARE-Luc I_max_ and anti-proliferative effects, confirming that the observed anti-proliferative activity is driven by NRF2 depletion (**Fig. 2J**). A set of cell lines (n=14) was chosen for a more comprehensive analysis of VVD-065 impact on proliferation. In 3D spheroid growth conditions, these cell lines showed a varying level of NRF2 dependency when NRF2 was genetically depleted (**Fig. S3H**). A subset of these NRF2-dependent cell lines demonstrated sensitivity to VVD-065 (**Fig. S3I**). NRF2 expression analysis revealed that all 4 responsive lines (KYSE70, NCI-H1793, HCC95, and LN18) are mechanistically sensitive (show loss of NRF2 protein expression) to VVD-065, whereas NRF2-dependent, but VVD-065-insensitive cell lines (NCI-H2122, NCI-H2023, A549, NCI-H1792) tended to show negligible changes in NRF2 protein levels (<10% reduction) (**Fig. 2K**). Intriguingly, these VVD-065-insensitive cell lines all harbor KEAP1 mutations (**Fig. S3H**) raising the possibility that many KEAP1 mutants do not bind NRF2 with sufficient residual affinity to support VVD-065-mediated NRF2 degradation (*33*).

### VVD-065 exhibits anti-tumor effects *in vivo*

Following oral dosing of VVD-065 in immunocompromised mice bearing KYSE70 tumor xenografts, plasma exposures increased dose-dependently, leading to robust intra-tumoral KEAP1_C151 engagement (**Fig. 3A**) and decreases in the expression of canonical NRF2 target genes (**Fig. 3B**). Robust NRF2 degradation was confirmed at the lowest dose level of 5 mg/kg (**Fig. 3C**). A time-course study in KYSE70 tumors revealed that covalent engagement of KEAP1_C151 by VVD-065 reaches maximal levels shortly after a single 5 mg/kg dose (**Fig. 3D**) and steadily declines over 3 days, at a rate consistent with a combination of compound clearance and KEAP1 protein half-life (11 hours). Repeat daily dosing of VVD-065 for 7 days resulted in higher levels of maximal engagement when compared to single administration consistent with incomplete protein turnover at 24-hours after a single dose. Reduced expression of NRF2 targets both at the transcript and protein level was observed following repeat dosing of VVD-065 (**Fig. 3D & S4A-C**).

**Figure 3:**
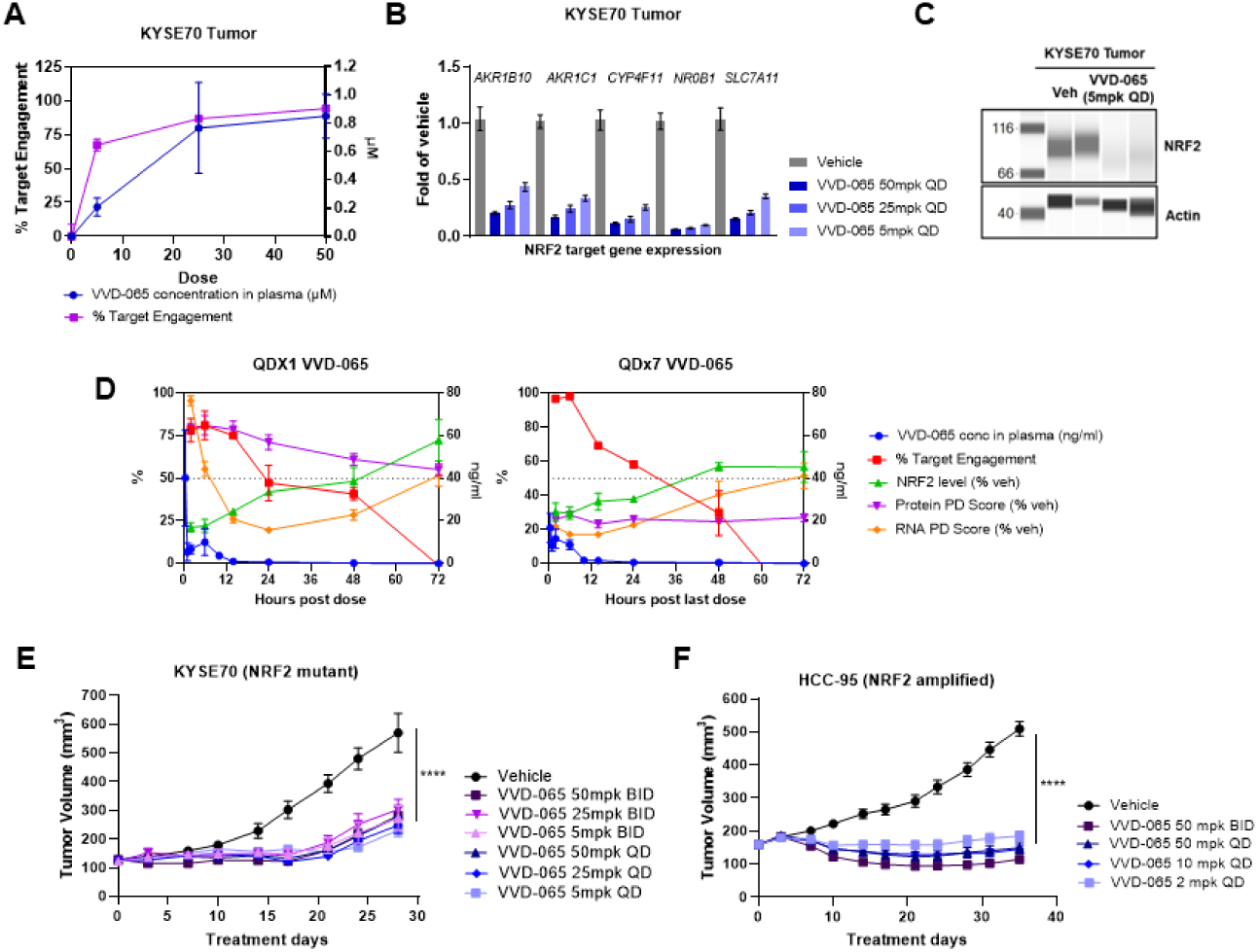
VVD-065 exhibits anti-tumorigenic effects *in vivo*. (A)(B) and (C) Pharmacokinetic (PK) and pharmacodynamic (PD) properties of VVD-065 in a KYSE70 xenograft model. KYSE70 tumor bearing animals were orally dosed at 5, 25, and 50 mg/kg once (A, B) or for 7 days (C). (A) Plasma was collected 30 minutes post dose for PK analysis and tumors were collected 24 hrs. post dose to measure covalent binding of VVD-065 to KEAP1_C151. (B) Expression of NRF2 target genes in KYSE70 tumors treated with VVD-065 at indicated doses measured by quantitative PCR. (C) Expression of NRF2 protein in vehicle or VVD-065 treated tumors. (D) PK - PD time course analysis in KYSE70 xenograft model. KYSE70 tumor bearing animals were orally dosed once (QDX1) or for 7 days (QDX7) with VVD-065 at 5mg/kg. Plasma exposure, KEAP1_C151 target engagement, NRF2 protein expression, and expression of NRF2 targets at RNA (AKR1B10, AKR1C1, ALDH3A1, CYP4F11, GPX2, NROB1, and SLC7A11) and protein (AKR1B10, AKR1C1, ALDH3A1, CYP4F11, and NROB1) levels were measured at indicated time-points (E) Anti-tumor efficacy of VVD-065 in KYSE70 xenograft model. Data are shown as mean ± s.e.m.; n=9-10 animals/ group. Mice were dosed orally with VVD-065 at indicated doses. (F) Anti-tumor efficacy of VVD-065 in HCC95 xenograft model. Data are shown as mean ± s.e.m.; n=10 animals/ group. Mice were dosed orally with VVD-065 at indicated doses. For figures E-F, statistical significance was calculated by 2-way ANOVA (ns, P > 0.05; *, P ≤ 0.05; **, P ≤ 0.01; *** P ≤ 0.001; ****, P ≤ 0.0001)

Treatment of a KYSE70 xenograft mouse model with doses of VVD-065 ranging from 5 to 25 mg/kg was well tolerated and resulted in robust tumor growth inhibition (TGI) (**Fig. 3E & S4D**). Genetic depletion of NRF2 in KYSE70 resulted in similar pharmacodynamic and TGI effects (**Fig. S4E-F**). TGI studies in additional cell lines revealed that responses in the 3D spheroid assay were generally predictive of *in vivo* efficacy. HCC-95 (NRF2^Amp^ sqNSCLC-squamous non-small cell lung cancer) showed strong responses to VVD-065 treatment in both 3D-sphere and *in vivo* settings (**Fig. 3F, S3I & S4G**) whereas KYSE180 (NRF2^D77V^; KEAP1^P278Q^ ESCC) and NCI-H23 (KEAP1^Q193H^ LUAD-lung adenocarcinoma) did not show growth inhibition in either setting (**Fig. S3I & S4H-K**) despite exhibiting pharmacodynamic (PD) responses to VVD-065 (**Fig. S4L**).

We expanded the *in vivo* evaluation of VVD-065 to genetically and histologically diverse patient-derived xenografts (PDX) (n=109), with a focus on sqNSCLC, ESCC, HNSCC (head & neck squamous cell carcinoma), and LUAD PDXs given the high-frequency of pathway mutations in these cancer types (*34–38*) (**Fig. 4A-D**). TGI responses were seen across both WT and KEAP1/CUL3/NRF2 mutant models, with the highest responses being observed in tumors enriched for mutations in the NRF2 pathway. A subset of PDXs were analyzed for pharmacodynamic responses, which identified three distinct groups– I) tumors that exhibited both PD responses and TGI following treatment with VVD-065 (**Fig. 4E & S5A-D**), II) tumors that did not show PD responses or TGI (**Fig. 4F & S5E**) and III) tumors that showed significant VVD-065-dependent NRF2 degradation, but with no impact on growth (**Fig. 4G & S5F-G**). These observations are consistent with non-adherent cell growth characterization of tumor lines, where models varied in both their intrinsic NRF2 dependence as well as their pharmacodynamic response to VVD-065. In a model where a strong response (partial regression) to VVD-065 was observed, the response was durable for at least 90 days with continued treatment (**Fig. 4H & S5H**).

**Figure 4:**
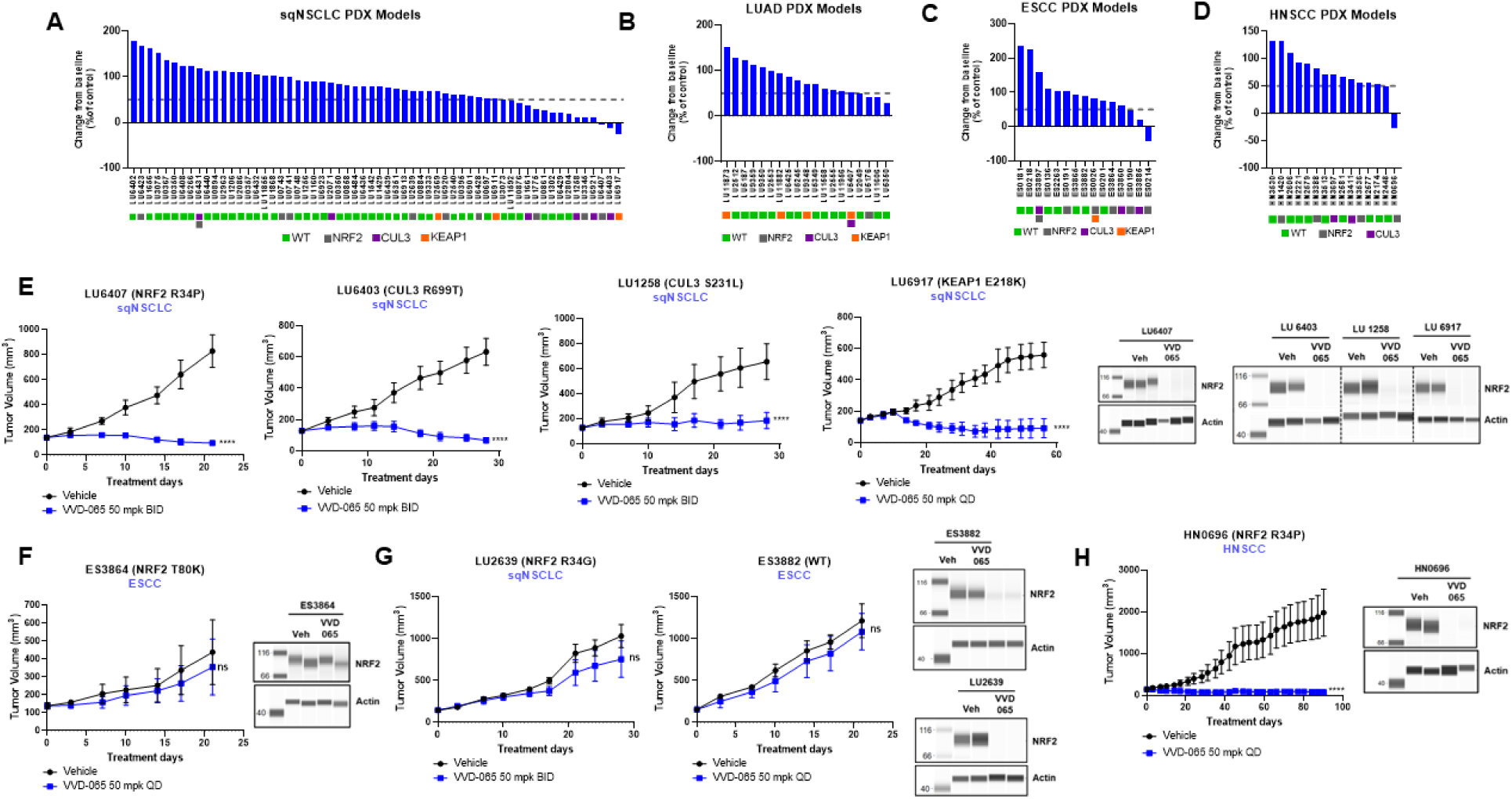
VVD-065 exhibits anti-tumorigenic effects in patient-derived xenograft (PDX) models. (A-D) Percent change in tumor volume on response calling day (days 20-28 after treatment initiation) in indicated patient-derived xenograft (PDX) models after VVD-065 administration. VVD-065 was administered orally at 50mg/kg QD or BID to mice bearing the indicated PDX models (n=2-3/ group). Bottom: amino acid changes in NRF2/ KEAP1/ CUL3 are shown for PDX models. Wild-type (WT) models are indicated by green filled boxes. (A) squamous non-small cell lung cancer (sqNSCLC), (B) lung adenocarcinoma (LUAD), (C) esophageal squamous cell carcinoma (ESCC) and (D) head and neck squamous cell carcinoma (HNSCC) PDX models. Dotted line represents 50% TGI threshold (E) Left: Anti-tumor efficacy of VVD-065 in LU6407, LU6403, LU1258, and LU6917 PDX models. Data are shown as mean ± s.e.m.; n=2-3 animals/ group. Mice were dosed orally with VVD-065 at indicated doses. Right: analysis of NRF2 protein expression in tumors from TGI study (F) Left: Anti-tumor efficacy of VVD-065 in ES3864 PDX model. Data are shown as mean ± s.e.m.; n=2 animals/ group. Mice were dosed orally with VVD-065 at indicated dose. Right: analysis of NRF2 protein expression in tumors from TGI study (G) Left: Anti-tumor efficacy of VVD-065 in LU2639 and ES3882 PDX models. Data are shown as mean ± s.e.m.; n=2-3animals/ group. Mice were dosed orally with VVD-065 at indicated dose. Right: analysis of NRF2 protein expression in tumors from TGI study (H) Left: Anti-tumor efficacy of VVD-065 in HN0696 patient-derived xenograft model. Data are shown as mean ± s.e.m.; n=3 animals/ group. Mice were dosed orally with VVD-065 at indicated dose. Right: analysis of NRF2 protein expression in tumors from TGI study For figures E-H, statistical significance was calculated by 2-way ANOVA (ns, P > 0.05; *, P ≤ 0.05; **, P ≤ 0.01; *** P ≤ 0.001; ****, P ≤ 0.0001)

### VVD-065 exhibits strong antitumor effects in syngeneic orthotopic settings

We were intrigued by the relatively high rate of both cell lines and PDX models with NRF2 pathway mutations showing strong pharmacodynamic responses to VVD-065 in the absence of effects on growth. While it is not unusual for a subset of cell lines or tumors harboring known oncogenic drivers to show no dependence on that driver (*39–41*), we considered that the complexity of NRF2 biology may not be fully recapitulated by heterotopic xenografts in immune compromised mice. Given the high oxygenation of the microenvironment, lung tissues and tumors are particularly vulnerable to oxidative stress, potentially contributing to the high rates of NRF2 pathway activation observed in lung cancers (*1, 42*). To explore potential contributors to NRF2 dependence in a more histologically relevant context, we tested VVD-065 effects on the growth of KLN-205 (WT murine sqNSCLC) cell line introduced in a syngeneic orthotopic setting (*43*). Despite being insensitive to VVD-065 in a 3D sphere assay (**Fig.S6A**), VVD-065 treatment resulted in a potent anti-tumor benefit with a 37% increase in median overall survival compared to vehicle treated animals (**Fig. 5A-B & S6B-C**). Modest but significant benefit was seen in the syngeneic heterotopic setting as well (**Fig. 5C**).

**Figure 5:**
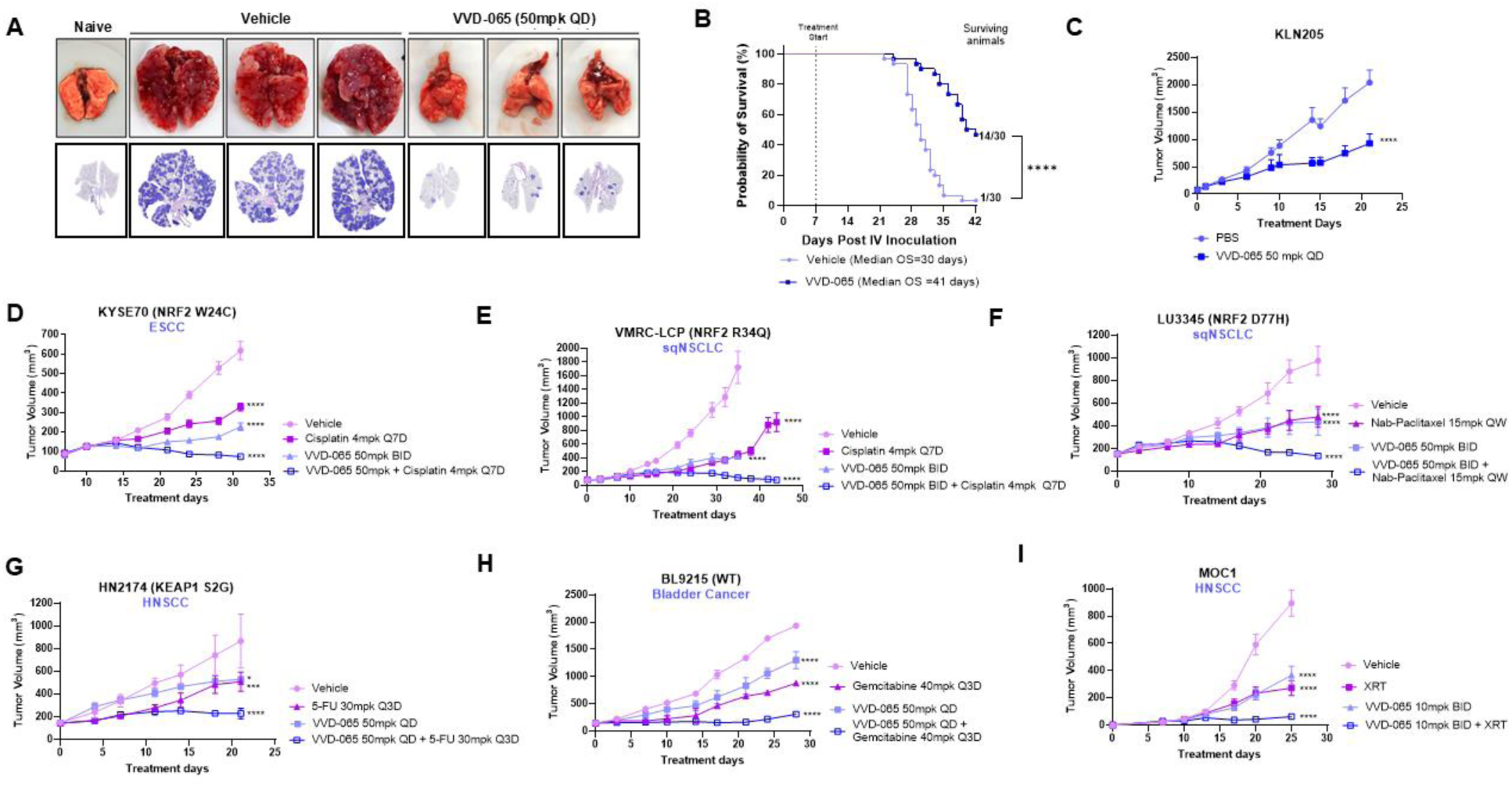
VVD-065 sensitizes tumors to chemo- and radiotherapy. (A) and (B) Anti-tumor efficacy of VVD-065 in KLN205 syngeneic orthotopic model. KLN-205 cells were injected into the tail vein of DBA/2 animals, and then dosed with vehicle or VVD-065 50mg/kg QD. (B) Representative images of lung from naïve animals, vehicle treated animals, and VVD-065 treated animals. (B) Kaplan-Meier survival analysis comparing VVD-065 to Vehicle (n=30 animals/ group) (C) Anti-tumor efficacy of VVD-065 in KLN205 syngeneic heterotopic model. Data are shown as mean ± s.e.m.; n=10 animals/ group. Mice were dosed orally with VVD-065 at indicated dose. (D) Anti-tumor efficacy of VVD-065 cisplatin combination in KYSE70 xenograft model. Data are shown as mean ± s.e.m.; n=9-15 animals/ group. Mice were dosed with VVD-065, cisplatin, or with combination of VVD-065 and cisplatin. VVD-065 was dosed orally, and cisplatin was dosed intra-peritoneally at indicated doses. (E) Anti-tumor efficacy of VVD-065 cisplatin combination in VMRC-LCP xenograft model. Data are shown as mean ± s.e.m.; n=10 animals/ group. Mice were dosed with VVD-065, cisplatin, or with combination of VVD-065 and cisplatin. VVD-065 was dosed orally, and cisplatin was dosed intra-peritoneally at indicated doses. (F) Anti-tumor efficacy of VVD-065 nab-paclitaxel combination in LU3345 PDX model. Data are shown as mean ± s.e.m.; n=3 animals/ group. Mice were dosed with VVD-065, nab-paclitaxel, or with combination of VVD-065 and nab-paclitaxel. VVD-065 was dosed orally, and nab-paclitaxel was dosed intra-venous at indicated doses. (G) Anti-tumor efficacy of VVD-065 in HN2174 PDX model. Data are shown as mean ± s.e.m.; n=2 animals/ group. Mice were dosed with VVD-065, 5-FU, or with combination of VVD-065 and 5-FU. VVD-065 was dosed orally, and 5-FU was dosed intra-peritoneally at indicated doses. (H) Anti-tumor efficacy of VVD-065 in BL9215 PDX model. Data are shown as mean ± s.e.m.; n=2 animals/ group. Mice were dosed with VVD-065, Gemcitabine, or with combination of VVD-065 and Gemcitabine. VVD-065 was dosed orally, and Gemcitabine was dosed intra-peritoneally at indicated doses (I) Anti-tumor efficacy of VVD-065 radiation combination at indicated doses in MOC1 syngeneic heterotopic model. Data are shown as mean ± s.e.m.; n=10 animals/ group. For figures D-I, statistical significance was calculated by 2-way ANOVA (*, P ≤ 0.05; **, P ≤ 0.01; *** P ≤ 0.001; ****, P ≤ 0.0001)

### VVD-065 shows combination benefit with chemo and radiotherapy

Since the upregulation of NRF2 is associated with chemo- and radio-resistance (*8, 44–48*), we tested VVD-065 in combination with these treatment modalities. In two squamous models (VMRC-LCP; sqNSCLC & KYSE70; ESCC), co-administration of VVD-065 and cisplatin exhibited a robust combination benefit without any significant additional weight loss (**Fig. 5D-E & S6D-E**). Co-administration of VVD-065 with other agents such as nab-paclitaxel, 5-FU, and gemcitabine also demonstrated robust benefits of combination therapy (**Fig. 5F-H & S6F-H**). Modest combination efficacy was observed with pemetrexed (**Fig. S6I-J**). Finally, we tested whether VVD-065 enhanced the radiosensitivity of tumors. In immunocompetent mice with subcutaneously implanted MOC1 (WT murine HNSCC) cells (*49*), VVD-065 or radiotherapy as monotherapy treatment achieved only modest TGI, whereas the combination of VVD-065 and radiotherapy showed nearly complete inhibition of tumor growth (**Fig. 5I**). These findings collectively indicate that pharmacological degradation of NRF2 can sensitize tumors to a range of chemo- and radiotherapies.

## Conclusion

The discovery of VVD-065 represents a new mechanistic paradigm for modulating the function of the KEAP1-NRF2 transcriptional circuit. By focusing primary screens on KEAP1_C151 engagement, followed by functional evaluation of confirmed binders, we discovered molecules with the unexpected ability to promote KEAP1-mediated NRF2 degradation through an allosteric molecular glue mechanism. These molecules enhance KEAP1 function by increasing KEAP1-CUL3 complexation, leading to enhanced NRF2 degradation in both WT and a significant subset of cancer-associated NRF2 pathway mutations. This contrasts with previous findings reporting KEAP1 inhibition through C151-reactive compounds and present a compelling and potentially generalizable case study for inducing degradation of a protein-of-interest through boosting the activity of its cognate E3 ligase. Further, we posit a new model wherein dynamic pharmacological regulation of the KEAP1-CUL3 interaction, as opposed to the KEAP1-NRF2 interaction, can govern NRF2 degradation.

While promising preclinical data suggest metabolic vulnerabilities of NRF2 hyperactivated tumors, clinical trials evaluating the glutaminase inhibitor CB-839 and mTOR1/2 inhibitor sapanisertib failed to demonstrate clinical benefits (*50*). A more effective approach entails direct targeting of NRF2. VVD-065 has demonstrated robust anti-tumor effects as a monotherapy and exhibited remarkable synergistic efficacy when combined with chemotherapy and radiotherapy. A related compound, VVD-130037, which employs the same molecular glue mechanism, is currently undergoing Phase 1 clinical trial (NCT05954312).

## Supporting information

PDB validation report

## Acknowledgement

The authors thank Benjamin F. Cravatt for reviewing the manuscript, Wuxi AppTec for compound synthesis, Novogene for RNA-Seq experiment, Synthego for CRISPR mediated knock-in cell line generation and Crown Biosciences and Champions Oncology for animal study support.

Studies conducted in the Major lab were supported by grants from the National Cancer Institute (T32-CA113275 Molecular Oncology Training Grant to I.B and CA216051 to M.B. Major).

The Berkeley Center for Structural Biology is supported by the Howard Hughes Medical Institute, Participating Research Team members, and the National Institutes of Health, National Institute of General Medical Sciences, ALS-ENABLE grant P30 GM124169. The Advanced Light Source is a Department of Energy Office of Science User Facility under Contract No. DE-AC02-05CH11231.

## Author contributions

N.R., T.M.K., M.P.P. conceptualized the project. N.R., J.P., R.T.A., T.M.K., G.M.S., G.A.E., M.B.M., D.S.W., and M.P.P. supervised the project. Investigation and experiments were carried out by T.W., I.C.L., J.M., C.L.E., S.M.B., I.B., A.N.S., A.P., J.C.G., J.I., S.R.J., B.K., N.I., S.G., H.P., D.B., S.S., P.Z., Z.R., K.N., H.N.W., E.W., M.K.P., J.J.S., E.T., and K.H. N.R. wrote the original draft. Writing, review, and editing were undertaken by N.R., J.C.G., G.M.S., M.B.M., D.S.W. and M.P.P.

---

**Supplemental Figure 1:**
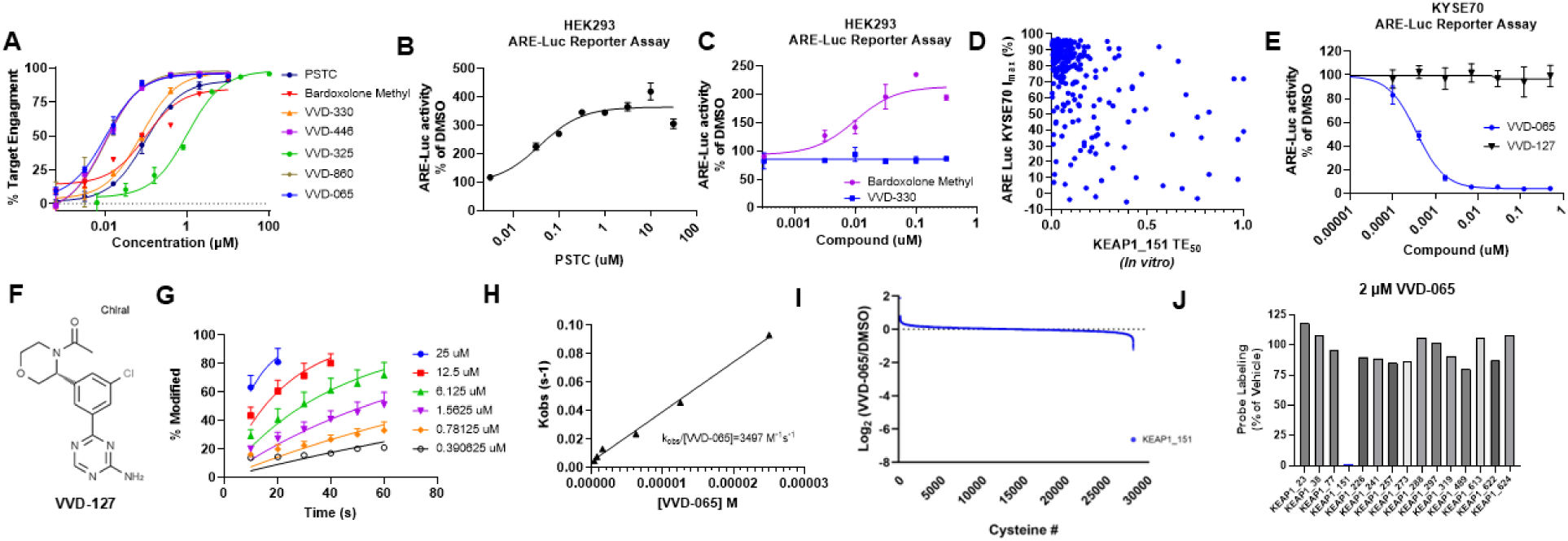
(A) Chemoproteomic determination of KEAP1_C151 engagement of indicated compounds. KYSE70 lysates were treated with DMSO or the indicated compounds. Exposed cysteines were then labeled by the addition of IA-DTB, and the samples were analyzed by PRM MS. Target engagement was calculated by comparing the area under the curve (AUC) for the KEAP1_C151 peptide of the VVD-065 treated samples to the AUC of the KEAP1_C151 peptide in the DMSO treated samples. (B) ARE-Luciferase reporter activity in HEK293 cells treated with PSTC (C) ARE-Luciferase reporter activity in HEK293 cells treated with Bardoxolone Methyl and VVD-330 (D) Correlation analysis of inhibition in KYSE70 ARE-Luc assay and covalent KEAP1_C151 target engagement for a set of KEAP1_151 compounds (E) ARE-Luciferase reporter activity in KYSE70 cells treated with VVD-127 (F) Structure of VVD-127, acetamide analog of VVD-065 (G) and (H) Reaction kinetics for VVD-065 determined using intact protein mass spectrometry with full length recombinant KEAP1. Target engagement measurements are the average of four replicates (I) Proteomic selectivity analysis of VVD-065 (>24000 sites were surveyed) (J) Analysis of covalent ligation of VVD-065 to different KEAP1 cysteines

**Supplemental Figure 2:**
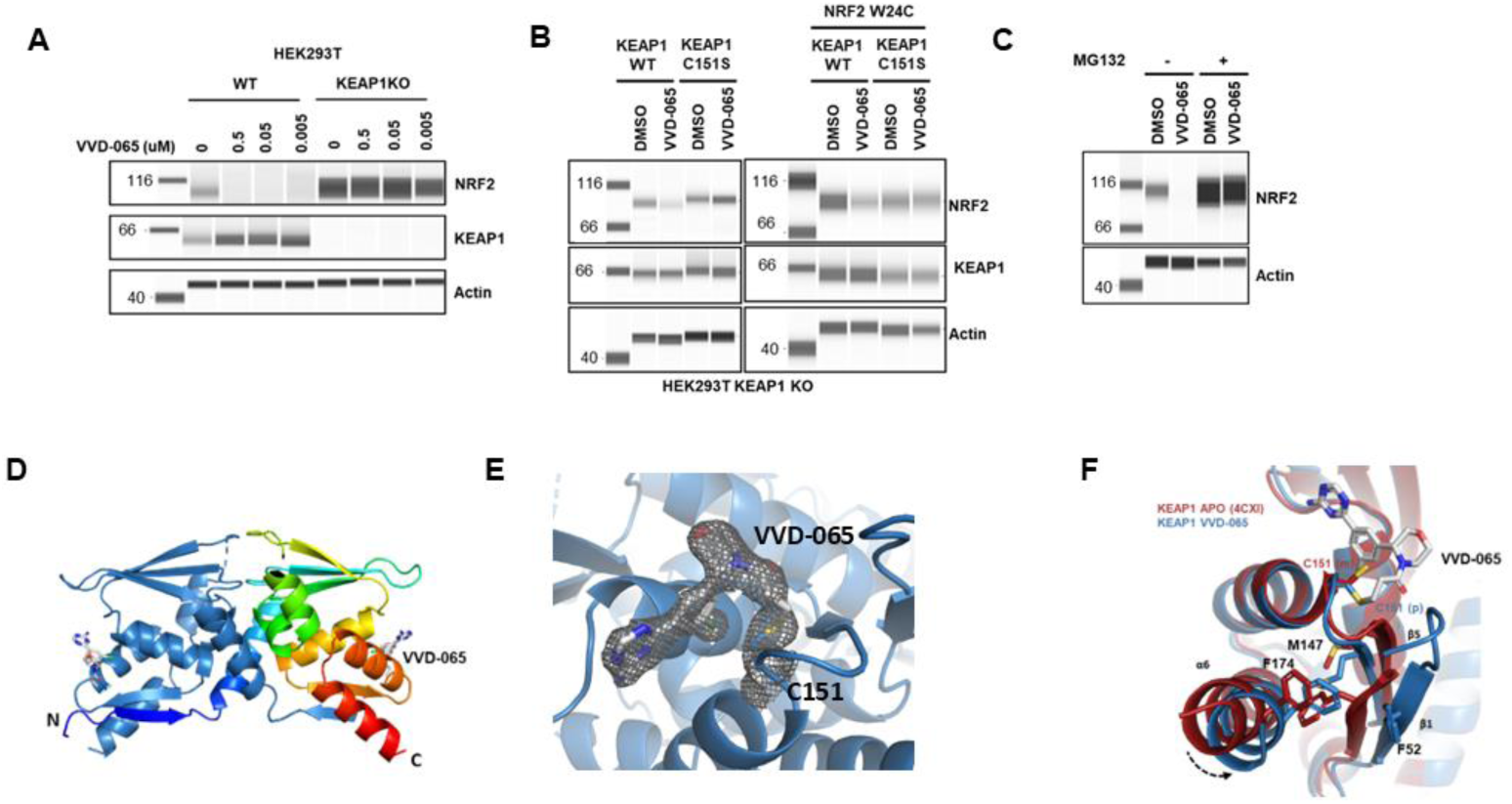
(A) Simple western analysis of NRF2, KEAP1, and Actin in HEK293T WT and HEK293T KEAP1 KO cells treated with VVD-065 for 18 hours (B) Left: Simple western analysis of VVD-065 mediated NRF2 degradation in HEK293T KEAP1 KO cells re-constituted with either WT KEAP1 or C151S KEAP1. Right: Simple western analysis of VVD-065 mediated NRF2 degradation in HEK293T KEAP1 KO cells overexpressing NRF2^W24C^ gain-of-function mutation and re-constituted with either WT KEAP1 or C151S KEAP1 (C) Simple western analysis of VVD-065 mediated NRF2 degradation in KYSE70 cells with or without MG132 co-treatment (D) Overall structure of the KEAP1 BTB domain bound to VVD-065 (E) 2F_O_-F_C_ simulated annealing omit map contoured at 1.5 σ showing clear electron density for VVD-065 continuous with KEAP1 C151. (F) Superposition of VVD-065 bound(blue) and APO KEAP1 BTB domain (dark red) highlighting the rotation of C151 and correlated movement of F52, M147 and F174

**Supplemental Figure 2:**
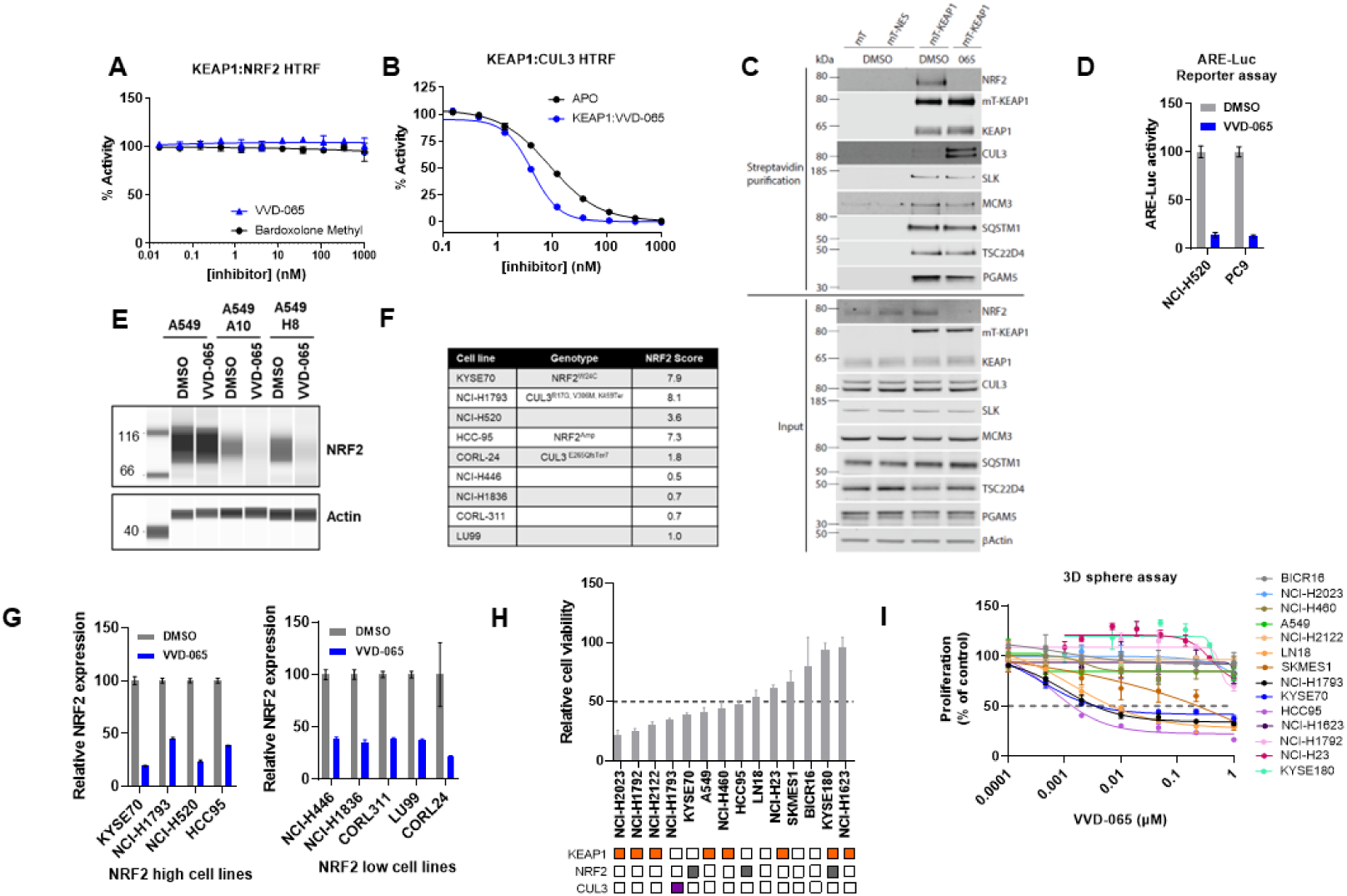
(A) Effect of Bardoxolone Methyl and VVD-065 on KEAP1 NRF2 interaction measured by HTRF assay (B) VVD-065 were pre-engaged with untagged KEAP1 construct and then compound activity on KEAP1 CUL3 interaction was measured by HTRF assay (C) HEK293 cells stably expressing N-termini miniTurbo tagged KEAP1 protein were treated with DMSO or VVD-065 for 2 hrs. and then incubated with biotin. Biotinylated proteins in cell lysate were enriched using streptavidin beads and analyzed via western blot using indicated antibodies (D) ARE-Luciferase reporter activity in PC9 and NCI-H520 cells (wild-type for KEAP, NRF2 and CUL3) treated for 18 hours with VVD-065. PC9 cells were treated with 0.2uM VVD-065 and NCI-H520 cells were treated with 0.3uM VVD-065 (E) Simple western analysis of NRF2 and Actin in A549 parental (KEAP1 G333C homozygous mutant) or A549 CRISPR edited cells (KEAP1 G333C heterozygous) treated with VVD-065. Two CRISPR edited knock-in clones (A10 and H8) were assessed (F) Genotype and NRF2 score of cell lines tested in Fig.S3G. NRF2 Score is geometric mean of AKR1B1, AKR1C1, ALDH3A1, CYP4F11, GCLC, GPX2, HMOX1, NQO1, SLC7A11, NROB1, and SRXN1 expression reported in CCLE. (G) NRF2 protein expression in VVD-065 treated NRF2 low and NRF2 high lines measured by ELISA (H) Viability levels of NRF2 intact (control) and NRF2-deficient (sgNRF2) cell lines in non-adherent 3D sphere assay. Bottom: Mutation status of NRF2/ KEAP1/ CUL3 are shown for each cell line. Filled boxes indicate presence of mutation (I) Viability levels of indicated cell lines treated with VVD-065 in non-adherent culture condition (3D sphere assay)

**Supplemental Figure 4:**
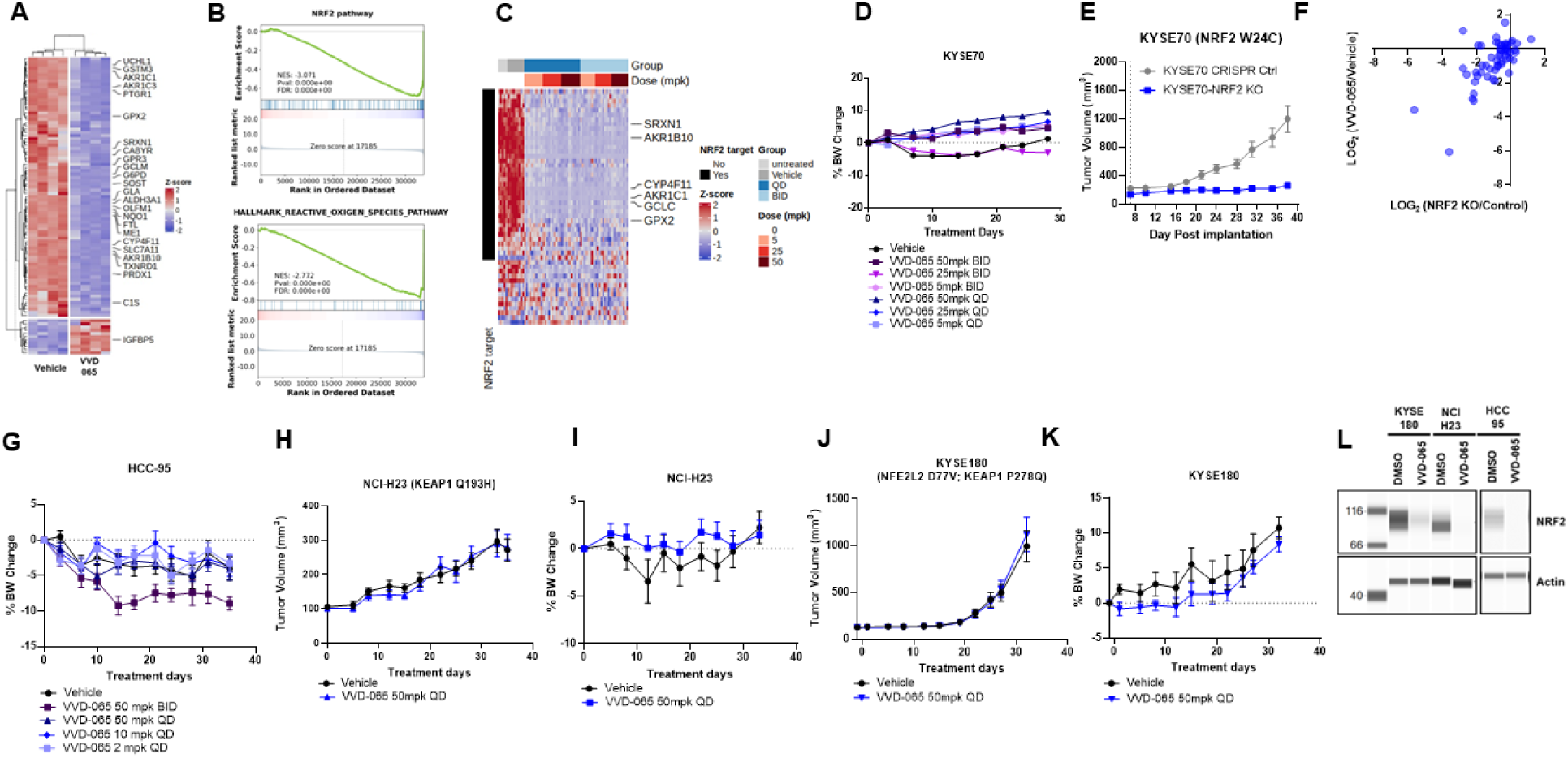
(A) , and (B) RNA-Seq analysis of KYSE70 tumors treated with either vehicle or VVD-065 at 5mg/kg for 7 days. (A) Heat-map of genes differentially expressed between vehicle and VVD-065 treated tumors. (B) GSEA analysis of differentially expressed genes between vehicle and VVD-065 treatment. (C) Analysis of NRF2 target expression at the protein level from KYSE70 tumor bearing animals dosed for 28 days. A set of proteins that are not canonical NRF2 targets was included in the analysis as a control. (D) Tolerability of VVD-065 in KYSE70 xenograft model. Data are shown as mean ± s.e.m.; n=9-10 animals/ group. Mice were dosed orally with VVD-065 at indicated doses. (E) Tumor growth kinetics of KYSE70 cells with or without NRF2 expression (CRISPR control vs NRF2 KO cells) (F) Correlation analysis of pharmacodynamic responses (expression of NRF2 targets at the protein level) between NRF2 KO tumors (vs. control) and VVD-065 treated tumors (vs. vehicle) (G) Tolerability of VVD-065 in HCC-95 xenograft models (H & J) Anti-tumor efficacy of VVD-065 in NCI-H23 & KYSE180 xenograft models. Data are shown as mean ± s.e.m.; n=10 animals/ group. Mice were dosed orally with VVD-065 at indicated doses. (I & K) Tolerability of VVD-065 in NCI-H23 & KYSE180 xenograft models (L) Analysis of NRF2 protein expression in HCC-95, NCI-H23, and KYSE180 treated with VVD-065 for 18 hrs.

**Supplemental Figure 5:**
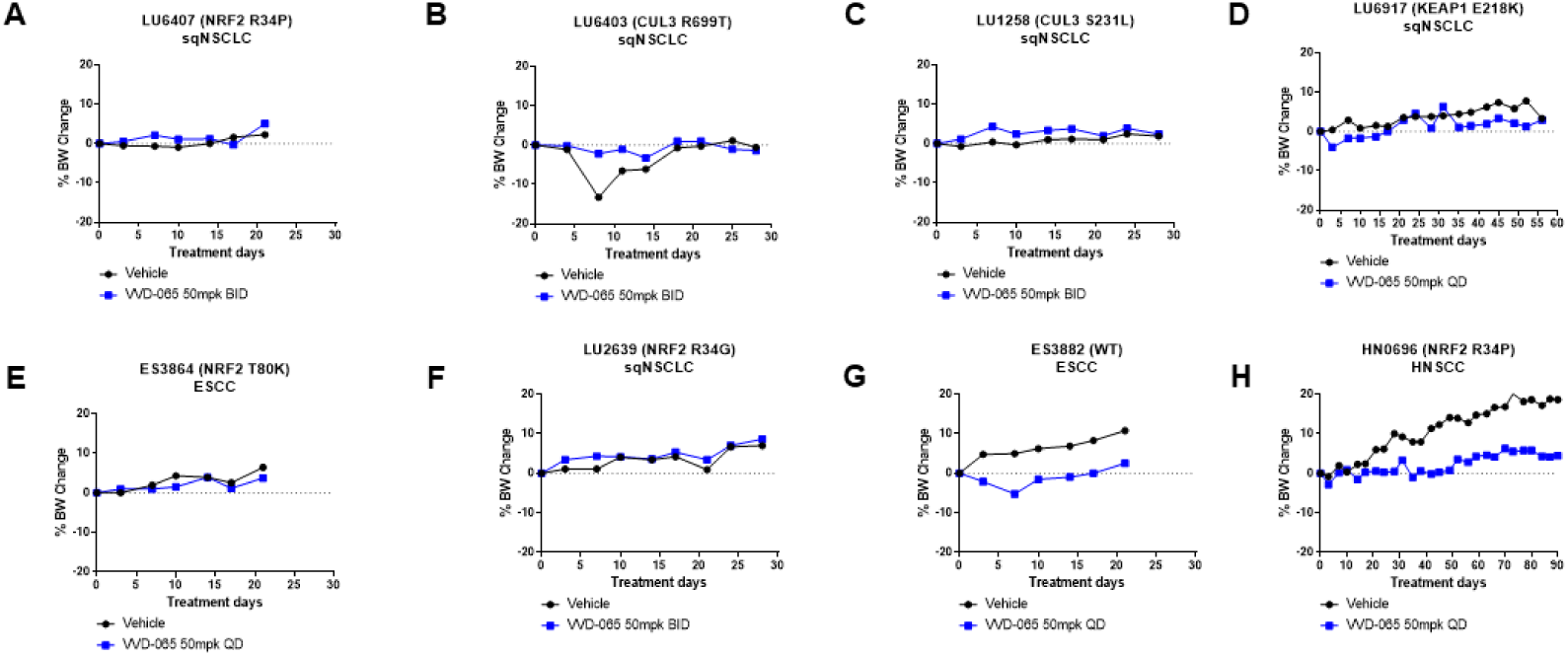
(A-H) Tolerability of VVD-065 treatment in models described in figure 4

**Supplemental Figure 6:**
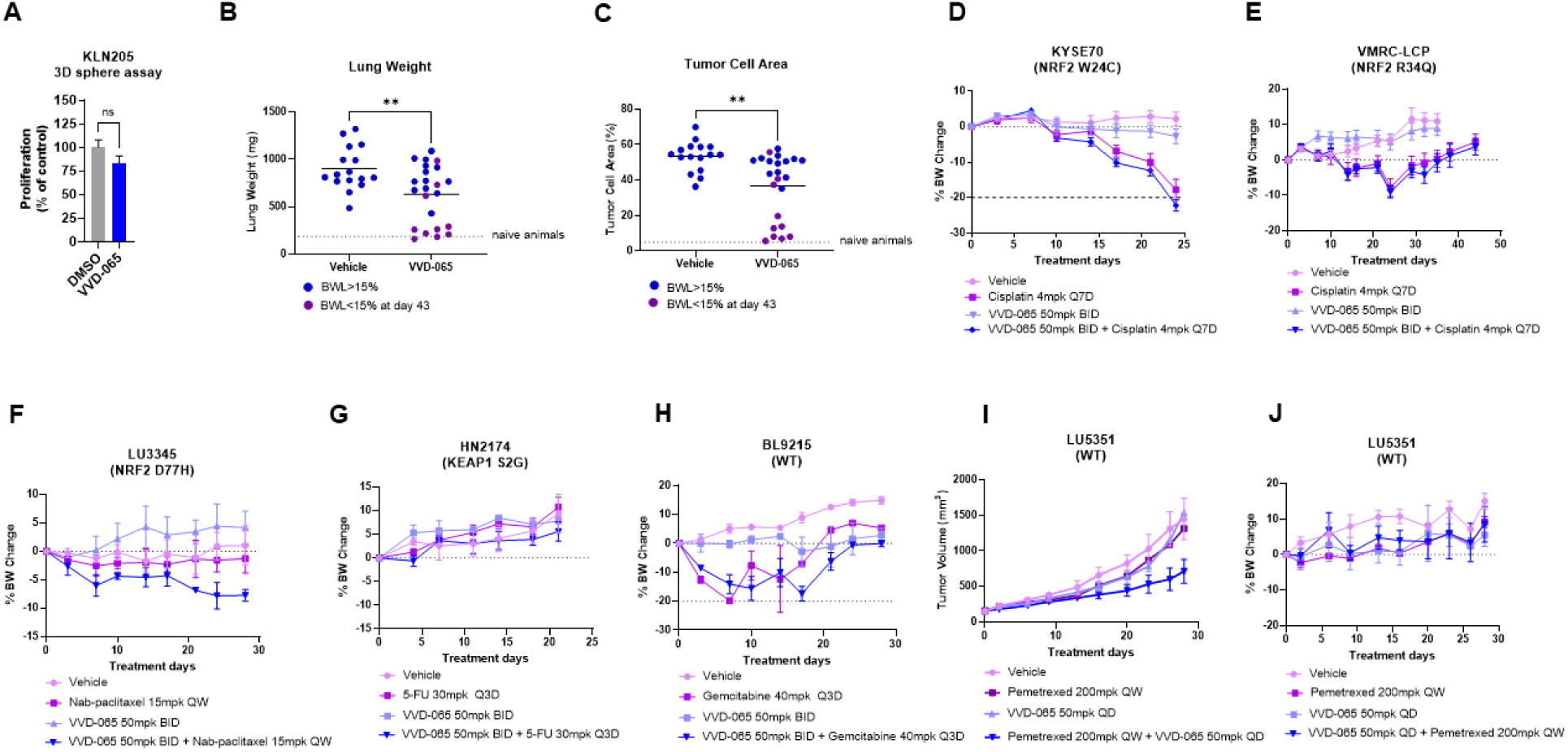
(A) Viability levels of KLN-205 cell line treated with 0.3uM VVD-065 in 3D sphere assay (B) and (C) Lung weight at termination and tumor cell area in vehicle or VVD-065 treated animals. Animals with body weight loss (BWL) of >15% are indicated by dark blue, and animals with BWL of <15% at day 43 are indicated by dark magenta (D-H & J) Tolerability of VVD-065 chemotherapy combination in indicated models (I) Anti-tumor efficacy of VVD-065 -/+ pemetrexed in LU5351 PDX model. Data are shown as mean ± s.e.m.; n=2 animals/ group. Mice were dosed with VVD-065, Pemetrexed, or with combination of VVD-065 and Pemetrexed. VVD-065 was dosed orally, and Pemetrexed was dosed intra-peritoneally at indicated doses

**Supplemental Table 2.** 1. Values in parenthesis include only reflections from the highest resolution shell.

| KEAP1 BTB domain in complex with VVD-065<br>PDB ID 9DU7 |  |
| --- | --- |
| Data collection |  |
| Wavelength (Å) | 1.00003 |
| Resolution (Å) | 48.14-1.87 (1.94-1.87)<sup>1</sup> |
| Space group | P4<sub>3</sub> |
| Unit cell (Å) | 72.7, 72.7, 64.2 |
| Unique reflections | 27282 (2674) |
| Multiplicity | 6.9 (7.2) |
| Completeness | 98.2 (97.0) |
| Mean I/sigma(I) | 15.2 (1.1) |
| Wilson B-factor | 42.6 |
| Rsym | 0.063 (1.34) |
| Rpim | 0.027 (0.54) |
| CC1/2 | 0.998 (0.525) |
| Refinement |  |
| Reflections using in refinement | 27280 (2674) |
| Reflections used for Rfree | 1332 (111) |
| R-work | 0.203 |
| R-free | 0.247 |
| RMS (bonds, Å) | 0.015 |
| RMS (angles, °) | 1.91 |
| Ramachandran allowed (%) | 99.19 |
| Ramachandran outliers (%) | 0.81 |
| Average B-factor | 47.6 |
| Protein | 47.0 |
| Ligand | 60.0 |
| Water | 52.2 |

## Materials and Methods

### ARE Luciferase reporter assay

KYSE70, LK2, EBC1, NCI-H520, PC9, NCI-H2228 and HEK293T cells were engineered with PiggyBac transposon system to contain NRF2-dependent ARE-Luciferase reporter cassettes. KYSE70, LK2, EBC1, NCI-H520, PC9, and NCI-H2228 cells were plated on Corning 96-well white/clear flat bottom microplates at 25,000 cells/well in 100 μL of appropriate cell culture media on Day 0. HEK293T cells were plated on Corning 384-well white, clear bottom microplates at 20,000 cells/well in 50uL media. All cells were incubated overnight (for a minimum of 18 hours) at 37°C and 5% CO^2^. On Day 1, cells were treated with compound. 6 hours later, to compound-treated HEK293T cells only, 50uL of reconstituted room temperature One-Glo EX reagent was added to each well and the contents were mixed for 2 minutes on a plate shaker at <200 rpm to lyse cells. Plates were then incubated for 3-5 minutes at room temperature to allow for stabilization of luciferase signal prior to measuring and reading Firefly luciferase signal on a Clariostar Plus plate reader. On Day 2, after 18-20 hours of treatment time of KYSE70, LK2, EBC1, NCI-H520, PC9, and NCI-H2228 cells, 80 µL of reconstituted room temperature One-Glo EX reagent was added to each well and the contents were mixed for 2 minutes on a plate shaker at <200 rpm to lyse cells. Plates were incubated for 3-5 minutes at room temperature to allow for stabilization of luciferase signal prior to measuring and reading Firefly luciferase signal on a Clariostar Plus plate reader.

### Compound treatment

Cells were treated with a dose response of Vividion compounds in 0.1 % DMSO using an HP Digital Dispenser. Each concentration was tested in duplicate.

For in vivo studies, VVD-065 was formulated in 10% NMP/90% Labrafil m1944CS.

For MG-132 experiment, cells were plated on Corning 96-well white/clear flat bottom microplates at 25,000 cells/well in 100 µl media on Day 0 and incubated overnight (for a minimum of 18 hours) at 37°C and 5% CO^2^. On Day 1, cells were pre-treated with 10uM MG132 for 5 hours prior to the addition of compound for 1 hour prior to harvesting for NRF2 expression analysis by Western blot.

### Protein Simple western blot analysis

#### In vitro

Cells were harvested for protein analysis by the addition of 25 μL of prepared lysis buffer (RIPA lysis buffer, Halt protease/phosphatase inhibitor cocktail, and benzonase nuclease; 25-29 U/μL) directly to cell culture plate wells. 8 μL of each cell lysate was mixed with 2 μL of 5X Jess sample buffer (provided in EZ Standard Pack and prepared as per manufacturer’s instructions). 10 μL Jess samples were boiled for 10 minutes at 95°C. The remaining lysate samples were used to determine the protein concentration using the Bio-Rad RC DC protein assay. After boiling, using the determined protein concentration, Jess lysates were normalized using 1X Jess sample buffer (5X diluted with prepared RIPA lysis buffer) to a final concentration of 1.0 μg/μL. 3 μL of each normalized lysate was loaded into each sample well on a pre-filled plate provided in the Jess Separation module; all corresponding reagent wells were loaded as per the manufacturer’s instructions, following the Replex protocol. The loaded plate was then centrifuged at 1000xg for 5 minutes to bring all contents to the bottom of each well. Both the sample loaded plate and a capillary cassette were loaded onto the Protein Simple machine as per the manufacturer’s instructions.

#### In vivo

One ml of cold Pierce IP lysis buffer (Thermo Scientific, PN 87788) was added to frozen tissues in bead beater tubes and tissues were homogenized by bead beating (Omni Bead Ruptor Elite) at 4°C for 30 seconds. Homogenized samples were centrifuged for 3 min at 7.4k x g. Supernatants were transferred to cluster tubes (Axygen, PN MTS-11-12-C) and treated with 24 µL of 1:1 solution of Benzonase (Millipore Sigma, PN 70746-3) and 100 mM MgSO4 (Sigma, PN M2643-500g) for 10 min at room temperature with shaking at 600 RPM. Lysates were cleared by centrifugation at 1000 x g, 4°C for 10 min and the supernatants were transferred to a 96 well plate (Lab Force, PN 1149J81). 8 µL of tissue homogenate was mixed with 2 µL of 5X Jess sample buffer (provided in EZ Standard Pack and prepared as per manufacturer’s instructions). 10 µL Jess samples were boiled for 10 minutes at 95°C. The remaining lysate samples were used to determine the protein concentration using Bio-Rad’s RC DC protein assay (catalogue no. # 5000122). Rest of the protocol follows the in vitro method described above.

For data analysis, raw chemiluminescence detection data was exported from the Jess instrument within the Compass for Simple Western software.

The following antibodies were used -

NRF2 (1:100 dilution – Cell Signaling Technology, # 20733)

β-actin (1:50 dilution – Cell Signaling Technology, # 4970)

KEAP1 (1:50 dilution – Cell Signaling Technology, # 8047)

### Cell lines and culture

Cell line source:

- KYSE70 cells (DSMZ; catalogue no. # ACC363)
- HCC95 cells (Sigma; catalogue no. # SCC483)
- VMRC-LCP cells (JCRB; catalogue no. # JCRB0103)
- NCI-H520 cells (ATCC; catalogue no. # HTB-182)
- NCI-H1793 cells (ATCC; catalogue no. # CRL-5896)
- EBC-1 cells (Accegen; catalogue no. # ABL-TC0170)
- NCI-H1623 cells (ATCC; catalogue no. # CRL-5881)
- NCI-H2228 (ATCC; catalogue no. # CRL-5935)
- LK-2 (RIKEN; catalogue no. # RCB1970)
- NCI-H23 (ATCC; catalogue no.# CRL-5800)
- KYSE180 (DSMZ; catalogue no.# ACC 379)
- SK-MES-1 (ATCC; catalogue no. # HTB-58)
- LN-18 (ATCC; catalogue no. # CRL-2610)
- NCI-H2122 (ATCC; catalogue no. # CRL-5985)
- A549 (ATCC; catalogue no. # CCL-185)
- NCI-H460 (ATCC; catalogue no. # HTB-177)
- NCI-H2023 (ATCC; catalogue no. # CRL-5912)
- BICR16 (Sigma; catalogue no. # 06031001)
- PC9 (Sigma; catalogue no. # 90071810)
- KLN-205 (ATCC; catalogue no. # CRL-1453)

Cell culture conditions:

Cell lines were maintained according to the vendor handling instructions.

### CRISPR/Cas9 mediated knock-in of KEAP1 C333G in A549

A549 has homozygous KEAP1 G333C mutation. Using proprietary methods, Synthego generated a knock-in cell line, where both mutant KEAP1 alleles were reverted to wild-type.

### CRISPR/Cas9 mediated knockout of NRF2 in cell lines

NFE2L2 Gene Knockout Kit was purchased from Editco Bio. containing guides: #1 GCGACGGAAAGAGUAUGAGC, guide_#2 AUUUGAUUGACAUACUUUGG, guide_#3 UAGUUGUAACUGAGCGAAAA. Guide pool was reconstituted using Nuclease-Free Duplex Buffer (IDT, # 11-01-03-01) to a final conc. of 100uM. NRF2 was knocked down using CRISPR editing via RNP electroporation. First, the RNP was complexed by mixing 3uL of Synthego NRF2 gene knockdown Kit V2 (pooled sgRNAs) with 2uL Cas9 (IDT; # 1081059). The complex was flicked to mix and incubated for 10-20 minutes at room temperature. While the RNP incubated, 500,000 cells were pelleted at 5K RPM for 1 minute, resuspended and washed in 1mL dPBS, and re-pelleted at 5K RPM for 1 minute. Cell pellets were resuspended in 20uL prepared electroporation buffer (16.4uL SE or P3 + 3.6uL supplement buffer per reaction) provided by the Lonza 4D-Nucleofector X kit (SE, Fisher Scientific, #NC0828118; or P3, Lonza; #V4XP-3032.) 20uL of the cell solution was then transferred to the 5uL RNP solution and gently mixed by pipetting. The cell + RNP mixture was then transferred to an 8-strip electroporation cuvette. Cells were electroporated using the CM137 setting. 100uL of warm serum-containing media was added to the cuvette to immediately recover cells. Nucleofected cells were immediately transferred to a 6 well plate containing 2ml of prewarmed media.

### Q-PCR analysis

#### In vitro

cells were processed for qPCR by 1) lysis in 50 μl/well of the Cells-to-CT kit provided lysis buffer, and 2) the addition of 5 μL STOP solution, as per the manufacturer’s instructions. 2 μL of each prepared cDNA lysate was then used directly for qPCR analysis by mixing with 1 μL of each PrimeTime™ assay probe, 5 μL TaqMan 1-Step qRT-PCR master mix, and 12 μL nuclease free water for a total 20 μL reaction mix. Prepared qPCR plate(s) were sealed with an adhesive cover before vortexing to mix samples, and a brief centrifugation to bring the well contents to the bottom of each well. The standard RT-qPCR cycling, as outlined in the manufacturer’s instructions were followed. For mRNA expression analysis, raw amplification data was exported into Microsoft Excel from the Quant Studio 6 instrument via the Thermo Fisher Connect Cloud functionality. Then, employing a delta Ct (ΔCt) method of analysis, ΔCt values for each sample were first calculated using housekeeping genes. Since all calculations are in the logarithm base 2, expression fold change was next calculated using the following formula: 2^-ΔCt. Expression fold change values were then used to calculate % inhibition relative to DMSO treated samples.

#### In vivo

Tissue harvested from mice was immersed in 800 µL DNA/ RNA shield (Zymo research R1200-125). After mechanical homogenization, homogenized tissue was spun down at 10,000 x g for 3 minutes to pellet cell debris. 200uL of supernatant were transferred to 96-well deep-well plate. Next, 10 µL of proteinase K solution was added to each sample and then incubated at room temperature for 30 minutes. After RT incubation, 200 µL DNA/ RNA lysis buffer, 30 µL of homogenously suspended mag-beads (Zymo research PN: R2130/R2131), and 400 µL of 100% ethanol (Acros organics: 61509-0010) were added to each well sequentially. Then, RNA was eluted using the Kingfisher Flex instrument. RNA concentration was measured afterward using Nanodrop. For real-time RT–PCR analysis, total RNA was reverse-transcribed to cDNA using High-Capacity cDNA Reverse Transcription Kit (Thermo Fisher PN: 4368814) followed by real-time PCR using either PerfeCTa FastMix II Low ROX (Quantabio PN: 97066-004) or TaqMan™ Fast Advanced Master Mix (Applied Biosystems PN: 4444557) with primers as described below. A total of ten genes were probed for expression as technical duplicates - four housekeeping genes B2M, HPRT1, TBP, 18S **or** ActinB & eight NRF2 target genes AKR1B10, AKR1C1, ALDH3A1, CYP4F11, GPX2, NR0B1, NQO1 and SLC7A11. Expression of 18S or ActinB was used to normalize mRNA levels of the gene of interest (dCT). Duplicate dCT values were then averaged, followed by normalization to vehicle group to obtain 2-ddCT. The 2-ddCT value of each gene of interest was normalized to the averaged 2-ddCT of Actin, B2M, HRPT1 and TBP to obtain fold change.

Probe information:

| Species | Assay ID | Fluorophore | Gene | Vendor |
| --- | --- | --- | --- | --- |
| Human | Hs.PT.58v.18759587 | FAM/Zen/IBFQ | B2M | IDT |
| Human | Hs.PT.58v.45621572 | FAM/Zen/IBFQ | HPRT1 | IDT |
| Human | Hs.PT.58v.39858774 | FAM/Zen/IBFQ | TBP | IDT |
| Human | Hs.PT.58.4411636 | FAM/Zen/IBFQ | CYP4F11 | IDT |
| Human | Hs.PT. 56a.24823646.g | FAM/Zen/IBFQ | ALDH3A1 | IDT |
| Human | Hs.PT.58.4044772 | FAM/Zen/IBFQ | SLC7A11 | IDT |
| Human | Hs.PT.58.26608447 | FAM/Zen/IBFQ | GPX2 | IDT |
| Human | Hs.PT.58.19137407 | FAM/Zen/IBFQ | AKR1B10 | IDT |
| Human | Hs.PT.58. 25746771.g | FAM/Zen/IBFQ | AKR1C1 | IDT |
| Human | Hs.PT.56a.1465041 | FAM/Zen/IBFQ | NR0B1 | IDT |
| Human | Hs99999901_s1 | VIC/MGB | 18s | Thermo |
| Human | 4326315E | VIC/MGB | ACTB | Thermo |
| Human | Hs.PT.58.2697277 | FAM/Zen/IBFQ | NQO1 | IDT |

### RNA-Seq analysis

RNA-seq experiment was conducted at Novogene. The gene expression omnibus accession number for the sequencing data reported in this manuscript is GSE278482.

### Plasmid information

KEAP1: WT KEAP1 (Promega; #N140A)

KEAP1 C151S: Q5 mutagenesis (NEB) performed *by* using Forward Primer “GGGCGAGAAGaGTGTCCTCCA” and Reverse Primer “ATGGAGATGGAGGCCGTG”

### Cell proliferation assay

#### 2D assay

Cells were plated on Corning 96-well white/clear flat bottom microplates at 500 cells/well in 100uL media on Day 0 and incubated overnight (for a minimum of 18 hours) at 37°C and 5% CO^2^. On Days 1 and 3, cells were treated with compounds. On day 5, 100uL of CTG reagent was added to all wells.

The contents were mixed for 2-3 minutes on a plate shaker and incubated for 10 minutes at room temperature to allow for stabilization of the CTG signal. Luminescence was then measured on the CLARIOstar plate reader.

#### 3D Sphere assay

Cells were plated on Corning 96-well black/clear round bottom microplates at 1000 cells/well in 100 µl media on Day 0 and incubated overnight (for a minimum of 18 hours) at 37°C and 5% CO2 to allow for 3D spheroid formation. On days 1,3,5,7, and 9, cells were treated with compounds. On Day 10, 100uL of CTG 3D reagent was added to all the wells. The contents were mixed for 10 minutes on a plate shaker, incubated for 25 minutes at room temperature to allow for stabilization of CTG signal.

Luminescence was then measured on the CLARIOstar plate reader.

#### Growth kinetics assay

For 2D culture 1,000 cells were added to each well of a 96 well plate (Corning® 96-well Flat Clear Bottom White Polystyrene TC-treated Microplates # 3610) in a volume of 200ul. For 3D culture 500 cells were added to each well of a 96 well spheroid plate (Corning® 96-well Spheroid Microplates # 4520) in a volume of 200ul. Cells were then grown for 7 days. On day 7 plates were imaged using the Incucyte SX5. Analysis was done with Incucyte software version 2022B Rev2. 2D analysis used Basic Analyzer Whole Well settings. Percent confluence was measured using an adjusted pixel size of -9 and min area (um2) 800. 3D analysis used Spheroid analysis settings. Largest Brightfield Object Area (um2) was determined with an adjusted pixel size of -10 and min area (um2) 8,000.

### In Vitro, In Situ and In Vivo Target Engagement

#### In vitro target engagement

Pellets derived from the indicated cell lines were lysed into DPBS by sonication and the protein concentration was determined via BioRad DC assay. Cell lysates were diluted to 2 mg/ml and treated with DMSO or compound for one hour at room temperature. Following incubation, IA-DTB was added to a final concentration of 200 µM and lysates were incubated for one hour at room temperature. 8x acetone was then added to the lysates and they were incubated at -80°C for two hours. Precipitated proteins were then pelleted by centrifugation (4,200 RPM, 45 min, 4°C).

#### In situ target engagement

For the indicated cell lines, cells were suspended at 2.5E^6^ cells/ml of RPMI, 10% FBS media. 1 ml of cell suspension was transferred to each well of a 96 well plate. Cells were treated with DMSO or compound and incubated in a 37°C CO_2_ incubator for two hours. After two hours, cells were pelleted (700xg, 10 min) and washed 2x with DPBS. After washing, cells were re-suspended in 200 µL DPBS and lysed by sonication using a QSonica water bath sonicator (50% amplitude, 30 sec on, 30 sec off, 2 mins total sonication). IA-DTB was added to 200 µM and lysates were incubated for one hour at room temperature. 8x acetone was then added to the lysates and they were incubated at -80°C for two hours. Precipitated proteins were then pelleted by centrifugation (4,200 RPM, 45 min, 4°C).

#### In vivo target engagement

Frozen tissues were homogenized by bead beating at 4°C for 30 seconds in IP lysis buffer (Thermo scientific). Homogenized tissues were centrifuged for 3 min at 7,400xg and the resulting supernatants were transferred to a 96 well plate and treated with 24 µL of a 1:1 solution of benzonase:100 mM MgSO_4_ for 10 min at room temperature. Lysates were then cleared by centrifugation at 1,000xg for 10 min at 4°C. Supernatants were transferred to a 96 well plate and protein concentration was determined via BioRad DC assay. Lysates were diluted to 2 mg/ml with DPBS and IA-DTB was added to a final concentration of 200 µM. Samples were then incubated with IA-DTB for one hour at room temperature. 8x acetone was then added to the lysates and they were incubated at -80°C for two hours. Precipitated proteins were then pelleted by centrifugation (4,200 RPM, 45 min, 4°C).

Protein pellets generated from in vitro, in situ and in vivo samples were further processed by resuspending the acetone pelleted material in 9M Urea, 50 mM ammonium bicarbonate. Free cysteine thiols were then reduced and alkylated by the addition of DTT and iodoacetamide (10 and 30 mM, respectively). After reduction and alkylation, samples were exchanged into 2M urea (Zeba spin desalting plates, Thermo Fisher) and digested with trypsin for 1 hour. IA-DTB labeled peptides were then isolated by the addition of 300 µL of 5% high-capacity streptavidin resin (Thermo Scientific). Bound peptides were washed 3x with wash buffer (0.1% NP-40, 150 mM NaCl, in 1x PBS) followed by four washes with HPLC grade water. Isolated peptides were eluted from streptavidin resin by the addition of 50% acetonitrile and dried in a speed vacuum concentrator.

### Target Engagement Quantification by Parallel Reaction Monitoring Mass Spectrometry

Isolated probe-labeled peptides were reconstituted in 3% acetonitrile, 0.1% formic acid. Peptides were concentrated onto an Acclaim PepMap100 C18 loading column (100 µm x 2 cm, 5 µm particle size, Thermo) and separated on a custom made C18 nanoviper analytical column (75 µm x 15 cm, 2 µm particle size, Thermo) using a Dionex Ultimate 3000 nano-LC (Thermo). Peptides were separated using a 12.7 min gradient from 6 to 32.5% solvent B (96.4% acetonitrile, 3.5% dimethylsulfoxide, 0.1% formic acid). Peptides were analyzed by parallel reaction monitoring (PRM) mass spectrometry using product ion scan mode on a Thermo Exploris 240 orbitrap mass spectrometer operated in positive ion mode. Detection of precursor ions was scheduled with 2 min acquisition windows. Product ion scan properties were as follows: 1.5 m/z Q1 resolution, normalized collision energy of 30%, 30,000 Oribitrap resolution, 70% RF lens, normalized AGC of 200% and dynamic injection time set to provide 7 points across a peak.

PRM data was analyzed using Skyline v.21.2.0.369 (MacCoss Lab, University of Washington). Peptide quantification was performed by summing the peak areas corresponding to six fragment ions pre-selected from an in-house reference spectral library. Retention Time Standard (RTS) peptides were used to normalize for sample variability. KEAP1 C151 target engagement is determined by monitoring the loss of IA-DTB probe labeling of the C151 containing peptide, CVLHVMNGAVMYQIDSVVR. Target engagement is estimated by comparing the average peptide area under the curve (AUC) for each dose group/test article to the average peptide AUC of the DMSO/vehicle group.

### Global chemoproteomics profiling

KYSE70 and MDA-MB-468 cell lysates were treated with 10, 2, 0.4, 0.08, 0.016, 0.0032 and 0.00064 µM VVD-8065 for one hour at room temperature and prepared as described for TE determination. After streptavidin enrichment and elution, the resulting dried peptides were resuspended in 70 µL of 0.2M EPPS pH 8.5, 30 µL of anhydrous acetonitrile and 55 µg of TMTpro 16plex reagent (Thermo, A44520).

After a two-hour incubation at room temperature, TMT labeling was quenched by the addition of 6.5µL of 5% hydroxylamine. Samples were then combined and desalted on a Biotage evolute express ABN plate. The resulting peptides were then separated into 96 fractions by high pH reversed phase chromatography. The 96 fractions were then recombined into 24 fractions and 12 of them where then subjected to MS analysis.

Peptides were reconstituted in 5% Acetonitrile, 0.1% FA and concentrated onto an Acclaim PepMap100 C18 loading column (100 µm x 2 cm, 5 µm particle size, Thermo) and then resolved on an Acclaim PepMap 100 analytical column (75 µm x 25 cm, 2 µm particle size, Thermo) on a Dionex Ultimate 3000 nano-LC (Thermo). Peptides were separated using a 171 min gradient from 6 to 30% solvent B (96.4% acentonitrile, 3.5% dimethylsulfoxide, 0.1% formic acid). The gradient then went from 30-40% B over one minute, 40-50% B over one minute and 50-60% B over one minuet. Quantitative TMT measurements were acquired on an Orbitrap Lumos Tribrid Mass Spectrometer (Thermo Scientific) using standard MS3 SPS data acquisition settings.

RAW files were searched with MassPike software package (GFY development team and the President and Fellows of Harvard University, GFY Core Version 3.4). Data was searched against the Human FASTA database with a peptide mass tolerance of 30 ppm, a fragment ion tolerance of 0.8, Trypsin digestion with a maximum of one internal cleavage site and a maximum of 3 differential modifications per peptide. Modification settings were as follows, static carboxyamidomethylation (+57.0214637236), static modification of Lysine and peptide N-terminus (+304.2071), differential oxidation of methionine (+15.99) differential IA-DTB labeling (+267.1945) of cysteines.

Searched data was exported from MassPike and further analyzed using our in-house data analysis algorithms. In brief, data is filtered such that all quantified peptides have an MS3 isolation specificity of ≥ 0.5, a sum signal for control wells of ≥ 40, control well CV < 0.5 and a max group standard deviation of < 25. Data is represented as % TE relative to controls. Data for all potential off-targets was manually inspected and sites were excluded from off-target consideration if replicate data was discordant, or TE did not exhibit a logical dose response.

### Crystallization, data collection and structure determination

Purified KEAP1 BTB domain was incubated with VVD-065 in 50 mM Tris pH 8.0, 150 mM NaCl, 1 mM TCEP, 2% DMSO and the reaction was monitored by intact protein MS. Upon completion, the protein sample was buffer exchanged into 20 mM Tris pH 8.0, 150 mM NaCl, 1 mM TCEP using a PD-10 desalting column and concentrated to ∼10 mg/ml. Crystals were grown in a 1:1 drop of protein to reservoir solution and equilibrated against 24% PEG 3350, 100 mM CHES pH 9.2 at 20 C. Crystals were cryoprotected by rapid transfer into reservoir solution supplemented with 25% glycerol before flash freezing in LN2. Diffraction data were collected on Advanced Light Source Beamline 5.0.2 and processed with XDS. The structure was determined by molecular replacement in Phaser using a single chain of 4CXT as a search model. The structure was refined using iterative rounds of refinement in REFMAC5 with manual inspection and model building in COOT. Ligand restraints for VVD-065 and the covalent bond with Cys151 were generated in JLigand. Waters were automatically added in COOT and REFMAC5 and manually inspected.

### Recombinant protein expression and purification

The gene for KEAP1 BTB domain for crystallization (residues 48-180 with the S172A mutation) was synthesized and cloned into pET28b with an N-terminal hexa-histidine tag and a thrombin cleavage site. The gene WT full length KEAP1 was synthesized and cloned into pFastBac1 with an N-terminal FLAG-6xHis tag followed by a TEV cleavage site. The gene encoding the N-terminal fragment of Cul3 (residues 1-388) was synthesized cloned into pGEX6P-1 with an N-terminal GST tag and Precision protease cleavage site. For E. coli expression, plasmids were transformed into BL21 DE3 Star. Large scale cultures were grown in 2XYT medium at 37C to an OD600 of 0.6 and induced with 0.4 mM IPTG at 18C overnight.

Full length KEAP1 was expressed in ExpiSf9 cells using baculovirus mediated expression in the EmBacY viral genome. KEAP1 BTB domain for crystallization was purified using the following procedure. Cells were resuspended in 50 mM HEPES pH 8, 500 mM NaCl, 10% glycerol, 10 mM imidazole and 1mM TCEP and lysed using a Microfluidizer. The lysate was cleared by centrifugation at 25,000 x g for 45 minutes before being loaded onto 5 ml of pre-equilibrated NiNTA resin. The resin was washed with 500 ml lysis buffer and eluted with lysis buffer supplemented with 250 mM imidazole. The 6xHis tag was removed with thrombin protease and the sample was dialyzed against lysis buffer. Uncleaved KEAP1 was removed with an additional NiNTA purification before being diluted 1:10 in 50 mM HEPES pH 8 and loading onto a POROS HQ IEX column. The protein was eluted with a 20 CV linear gradient from 50 mM to 1M NaCl and fractions containing KEAP1 were pooled, concentrated, and injected onto a Superdex S200 column equilibrated in 25mM HEPES, pH 8, 150mM NaCl, 1mM TCEP. Full length KEAP1 was purified in a similar manner, omitting the proteolytic cleavage and IEX steps.

### Intact protein mass spectrometry and rate determination

To observe covalent ligand engagement of recombinant KEAP1, samples were analyzed following formic acid quenching on an Agilent LC1290 Infinity II instrument coupled to a 6545 QTOF liquid chromatography–mass spectrometer (Agilent Technologies). A sample volume of 10 μl, equivalent to approximately 1.5 pmol of KEAP1 protein, was injected. The protein was desalted and separated on an AERIS 3.6-μM-wide-bore XB-C8 liquid chromatography column (50 × 2.1 mm^2^, Phenomenex) at 60 °C at a flow rate of 0.5 ml min^−1^. Liquid chromatography solvent A comprised 0.1% formic acid in 99.9% water, and solvent B was 0.1% formic acid in 99.9% acetonitrile. The column was equilibrated in 10% B for 30 s, followed by a 3.5 min gradient from 10 to 70% B to separate the analytes. This was followed by a 15 s gradient from 70 to 95% B, a 15 s gradient from 95 to 10% B, a 15 s gradient from 10 to 95% B and finally a 15 s gradient from 95 to 10% B to clean the column before re-equilibration. Mass spectra were acquired from 700 to 1,700 Da at a resolution of 25,000. A Dual Agilent Jet Stream Electrospray Ionization Source was used for ionization. The gas temperature was set to 325 °C, with a flow rate of 10 l min^−1^. The nebulizer was set to 45 pounds per square inch and sheath gas temperature and flow were set to 375 °C and 12 l min^−1^, respectively. One spectrum was acquired per second with a collision energy of 10 V. The capillary voltage was set to 5,000 V and the nozzle voltage to 2,000 V. The fragmenter, skimmer and octopole radio frequency (RF) peaks were set at 250, 65 and 750 V, respectively. The resulting data files were deconvoluted to protein masses using Agilent MassHunter BioConfirm Software, v.11.0. The biomolecule table containing protein mass and peak intensities was used to quantify the percentage of compound modification relative to the unmodified protein peak, by dividing modified protein intensity by the sum of the unmodified and modified protein intensities. *k*_obs_/[*I*] was calculated using assumptions for pseudo-first-order reaction kinetics (d(VVD-065)/d*t* = −*k* × (VVD-065), (VVD-065)*_t_* = (VVD-065)*_t_*_0_ × e^−*kt*^) and averaged for each inhibitor concentration; *k*_obs_ was determined by dividing these values by (*I*) for each concentration.

### KEAP1-CUL3 and KEAP1-NRF2 HTRF assays

#### KEAP1 Buffer

| [Initial] | Name | [Final] | Amount |
| --- | --- | --- | --- |
| 1M | TRIS pH8 | 50mM | 1mL |
| 1M | MgCl<sub>2</sub> | 5mM | 100uL |
| 5M | NaCl | 100mM | 400uL |
| 30% | BSA | 0.005% | 3.33uL |
| 1M | DTT | 1mM | 20uL |
| 100mM | CHAPS | 2mM | 400uL |
|  | H<sub>2</sub>O |  | 18mL |
|  |  | Total: | ~20mL |

HTRF Assay: KEAP1 diluted to 12 nM in KEAP1 buffer was plated at 5uL/well. Compound was added to plate via compound printer and incubate for 15 min at room temperature. Next, CUL3 diluted in KEAP1 buffer, 5uL/well was plated. Plate was spun and pulsed to 1000g, followed by incubation for 1hr. Donor/Acceptor solution (anti-His-Tb Gold and anti-GST-d2 both diluted 1:100 in PPI Tb detection buffer) was added at 10uL/well and incubated for 1 hr. Luminescence reading was measured in Clariostar. Same general protocol was used for KEAP1:NRF2 HTRF assay. Final concentrations of 2 nM FL FLAG-KEAP1 and 1 nM GST-NRF2 were used, and the dyes used were anti-FLAG-Tb and anti-GST-d2.

### NRF2 ELISA

Cells were seeded and treated as indicated with either DMSO or the indicated concentrations of VVD-065 for 18-24 hour at 37C. Following treatment cells were washed and lysed with minimal volumes of Pierce IP lysis buffer (+PIC, MgS_4_ and benzonase) and incubated for 1 hour at room temperature. Then lysate was diluted with 100 µL of dilution buffer to the indicated concentrations, centrifuged for 5 minutes at 4122 g and stored at -80°C.

The NRF2-specific capture antibody (Cell Signaling 12721S) was diluted in plating buffer and added to assay plate, 20 µl/well at 5 ng/µl, overnight at 4°C. The plate was washed 2 times in PBST and the blocking buffer was added, 100 µl/well for 1h at room temperature. Following incubation, the blocking buffer was removed, and lysate diluted in dilution buffer was added for 1.5 hours and room temperature, 25 µl/well. The assay plate was washed 3 times with 100 µl/well of PBST and 20 µl/well biotinylated detection antibody (Abcam, Ab62352) was diluted (1:100 in dilution buffer) and added for 1 hour at room temperature. The assay plate was washed 3 times with 100 µl/well of PBST and 20 µl/well Neutraviden-HRP diluted in dilution buffer was added for 1 hour at room temperature. The assay plate was washed 4 times with PBST, 100 µl/well. HRP substrate was added to plate, 20 µl/well, prior to the Clariostar luminescence read. Data were normalized to untreated wells (100%).

### RNA-Seq data analysis

All statistical analyses were done in python (3.9.15) and R (4.3.2). Differential expression was done using DESeq2 (1.40.2) and limma (3.56.2) (Ritchie ME et al., 2015). Gene set enrichment analysis was done using gseapy (0.14.0) (Subramanian A et al., 2005; Mootha VK et al., 2003). Visualizations were made using matplotlib (3.6.2), seaborn (0.13.2), and complex heatmap (2.16.0) (Gu et al., 2016; Gu Z., 2022). Principal component analysis was done using scikit-learn (1.1.3).

### Plasma VVD-065 measurement

VVD-065 was solvated in DMSO (ACS reagent >/=99.9%, Sigma Aldrich) to prepare 0.5 mg/mL stock, which was stored at -20°C until use. Reagents used for bioanalysis included acetonitrile and water (high-performance liquid chromatography grade, Fisher Scientific) and formic acid (Optima grade, Fisher Scientific), while consumables included 96-well polypropylene plates (Nunc-Thermo) and 96-well silicone plate maps (Axygen). Prior to bioanalysis, working solutions were serially prepared in 75:25 acetonitrile/water and spiked (20 µL) into commercially obtained mouse plasma containing K2-EDTA anticoagulant (BioIVT, Baltimore, MD) for preparation of standards ranging in concentration from 0.500 to 5000 ng/mL. Study samples were removed from frozen storage and thawed, where 20 µL plasma aliquots (2 µL aliquots combined with 18 µL of blank commercial mouse plasma for 10-fold dilutions and 10 µL aliquots combined with 10 µL of blank commercial mouse plasma for 2-fold dilutions) were transferred to clean 96-deep well plates for extraction by protein precipitation via addition of 200 µL 95:5 acetonitrile/methanol (high-performance liquid chromatography grade, Fisher Scientific) containing carbamazepine and verapamil internal standard followed by sealing, vortex mixing (11 minutes) and centrifugation (4121-G at +4°C for 10 minutes). For the final sample preparation step, 180 uL of supernatant was transferred to a clean 96-well plate, sealed with a silicone mat, and submitted for Liquid Chromatography with tandem Mass Spectrometry (LC-MS/MS) analysis. Upon injection of sample (9 µL), chromatographic separation was achieved using a Kinetex Biphenyl, 2.1×50 mm, 2.6-micron, 100A HPLC column (Phenomenex, Torrance, CA) maintained in a column oven at 39°C using a binary gradient 5% to 95% B, where pump A delivered water containing 0.1% formic acid and pump B delivered acetonitrile with 0.1% formic acid for a total flow rate of 0.65 mL/min and 2.1 minutes total run time. A linear ion trap mass spectrometer (Sciex 5500 or Sciex 6500+, Framingham, MA) was operated in either positive ion mode electrospray ionization or positive ion mode atmospheric pressure chemical ionization with detection by multiple reaction monitoring. The acquired raw data were processed using Analyst® software (version 1.7.1), where sample concentrations were determined using a linear calibration curve comprising a minimum of 8 nonzero standards and appropriate weighting. The minimum batch acceptance criteria included calibrator accuracy ±20%, correlation coefficient (r) >0.99 and 2/3 quality control samples within ±25% the theoretical value.

### In vivo studies

#### Human cell-line derived xenograft (CDX) studies

Following the acclimation period (3-7 days), immunodeficient mice, NSG (Jackson Laboratory) for KYSE70 and VMRC-LCP inoculation, and NCG (GemPharmatech Co, China) for HCC95 inoculation, were inoculated subcutaneously into the dorsal flank with a cell suspension of 90% viable tumor cells in 0.2 ml of serum-free RPMI-1640: HC Matrigel or regular Matrigel (Corning, catalogue no. # 354262; Corning, cat# 356237) (1:1 v/v) delivering tumor cells (KYSE70 and VMRC-LCP - 5 × 10^6^ cells/ mouse; HCC95 - 1X10^7^ cells/ mouse). Mice were randomized based on tumor volume and were enrolled into different groups. Tumor volumes were measured twice per week after randomization in two dimensions using a caliper, and the volume was determined using the formula: V = (L x W x W)/2, where V is tumor volume, L is tumor length (the longest tumor dimension), and W is tumor width (the longest tumor dimension perpendicular to L). Tumor growth inhibition was calculated using the formula: TGI%=[1-(Ti-T0)/(Ci-C0)]x100; Ti as the mean tumor volume of the treatment group on the measurement day, T0 as on the initiation day; Ci as the mean tumor volume of control group at the measurement day, C0 as on the initiation day. VVD-065 was administered at the indicated dose level by oral gavage. HCC95 CDX TGI was conducted at Crown Biosciences. Tumor tissue for ex vivo analysis was collected at 14 hours post final dose in a BID regimen and 24 hours post final dose in a QD regimen.

#### Mouse syngeneic TGI study

KLN-205 syngeneic TGI study was conducted at Champions Oncology. 2X10^5^ KLN-205 cells per mouse were subcutaneously implanted in DBA/2 mice (Jackson Laboratory). Rest of the study was conducted as described above.

#### Radiation combination study

MOC1 cells were generous provided by Dr. Ravindra Uppaluri (Dana Farber Cancer Institute, Boston, MA) and cultured in medium with 5% FCS, 40 ng/mL hydrocortisone (catalog #H0135, Sigma-Aldrich), 5 ng/mL EGF (catalog #01–107, Sigma-Aldrich), 5 mg/mL insulin (catalog #I0516, Sigma-Aldrich), and 1% penicillin–streptomycin. Eight to ten-week-old female C57BL/6 mice were purchased from Taconic Laboratory. A total of 5×10^6^ MOC1 cells were subcutaneously injected into the dorsal flanks of the mice. Tumor-bearing mice were randomized after 7 days to either vehicle control (Labrafil M 1944 CS 5%DMSO), VVD065 (10mg/kg bid) days 7-28, RT alone days 14-18, or VVD065+RT. RT was performed using the Small Animal Radiation Research Platform 200 (SARRP) from XStrahl (Suwannee, GA) for a tumor dose of 2Gy daily on 5 consecutive days (10Gy total).

#### Patient derived xenograft (PDX studies)

PDX studies were conducted at Crown Biosciences. Fresh tumor tissues from mice bearing established primary human cancer PDX model was harvested and cut into small pieces (approximately 2-3 mm in diameter). PDX tumor fragment, harvested from donor mice, was inoculated subcutaneously at the upper right dorsal flank into immunocompromised mice for tumor development. Mice were randomized based on tumor volume (100-200 mm3). After randomization, tumor bearing mice were allocated into 2 or 4 groups with 2-3 mice per group. Tumor tissue for ex vivo analysis was collected at 14 hours post final dose in a BID regimen and 24 hours post final dose in a QD regimen.

#### KLN-205 disseminated model

KLN-205 cells were harvested in exponential growth phase and suspended in 100 uL PBS to deliver 0.5×10^6^ cells via a tail vein injection in DBA/2 mice (Jackson Laboratory). 7 days after inoculation, mice were randomized based on initial body weight and indicated treatment was dosed QD via oral gavage. Health observations and body weights were recorded daily. Mice reached their endpoint upon body weight loss of 15% compared to treatment initiation, which was validated to capture mice prior to health deterioration. Upon euthanasia, lung weights were recorded, followed by inflation with 10% NBF for subsequent formalin-fixed paraffin-embedding. Tumor nodules were quantified on H&E-stained lung sections.

### Affinity purification of biotinylated proteins

HEK-293T cells stably expressing empty-miniTurbo (mT-V5) or miniTurbo-tagged nuclear export signal (mT-NES) or KEAP1 protein (mT-KEAP1) were treated with 100 nM VVD-065 (mT-KEAP1) or DMSO (mT-V5, mT-NES, mT-KEAP1) for 4 hours. Cells were then treated with 50 uM biotin (Sigma, Cat. # B4639-1G) for last 1 hour. Cells were washed with cold PBS and harvested on ice. Cells were sonicated at 4°C in RIPA buffer (10% glycerol, 50 mM HEPES, 150 mM NaCl, 2 mM EDTA, 0.1% SDS, 1% Triton X-100, 0.2% Sodium deoxycholate) containing protease inhibitor (Thermo Scientific, Cat. # 78429), phosphatase inhibitor (Thermo Scientific, Cat. # 78426), and benzonase (Sigma, Cat. # E1014-5KU). Protein lysates were incubated with 30 µL of packed pre-washed Streptavidin Sepharose beads (Cytiva, Cat. # 17511301) overnight on a rotator at 4°C. Beads were washed with wash buffer (WB) 1 (2% SDS) twice, once with WB2 (500 mM NaCl, 0.1% deoxycholate, 1% Triton X-100, 1 mM EDTA, 50 mM HEPES, pH 7.5), once with WB3 (250 mM LiCl, 0.5% Triton X-100, 0.5% deoxycholate, 1 mM EDTA, 50 mM HEPES. pH 8.1), and once with WB4 (150 mM NaCl, 50 mM HEPES, pH 7.4). After washing, biotinylated proteins were eluted in 2X LDS sample buffer (Invitrogen, Cat. # NP0007) with 2 mM biotin and 50 mM DTT (Sigma, Cat. # D0632-5G) for 10 minutes at 70°C.

### Immunoblotting

Proteins were separated in 4-12% Bis-Tris gel (Invitrogen, Cat. # NP0321BOX) using 1X MOPS buffer (Fisher/Boston Bioproducts, Cat. # NC0778891) and transferred to nitrocellulose membranes (Thermo Scientific, Cat. # PI88018). Membranes were blocked in 5% milk in TBST and incubated with primary antibodies diluted in 5% BSA overnight at 4°C. Membranes were then washed in TBST and incubated with secondary antibodies, 1:10,000 in 5% milk in TBST (Fisher/LI-COR, Cat. # NC0250903 and NC0250902) at room temperature for 1 hour. Imaging was performed in Odyssey CLx and analyzed in Image Studio Lite Ver 5.2 software. Primary antibodies used were as follows: NRF2 (Cell signaling, Cat. # 2073S, 1:1000), CUL3 (Cell signaling, Cat. # 2759, 1:1000), KEAP1 (Cell signaling, Cat. # 8047, 1:1000), SQSTM1 (Bethyl, Cat. # A302-856A-T, 1:1000), TSC22D4 (Bethyl, Cat. # A303-222A, 1:200), PGAM5 (Abcam, Cat. # ab126534, 1:1000), MCM3 (Bethyl, Cat. # A300-192A, 1:2000), SLK (Bethyl, Cat. # A300-500A, 1:1000) and β-Actin (Sigma, Cat. # A5316, 1:5000)

### LC-MS/MS sample preparation, data acquisition, raw data processing, and analysis

Sample preparation for mass spectrometry analysis: HEK-293T cells stably expressing N- and C-termini miniTurbo biotin ligase tagged KEAP1 were treated with 50 nM VVD-065 or DMSO for 4 hours and 50 µM biotin during the last one hour. Affinity purification of biotinylated proteins was performed as described earlier. For mass spectrometry, protein lysate collected from a 15 cm tissue culture plate were incubated with 30µL packed Streptavidin Sepharose beads. Beads were then washed in WB1-4 as described earlier. The streptavidin beads were further washed 3X in 50 mM ammonium bicarbonate (ABC). Beads were then resuspended in 100 µl of 50 mM ABC containing 1 µg trypsin (Promega, Cat. # V5113) and 0.1 mAu Lys-C (Wako Chemicals, Cat. # 129-02541) and incubated overnight at 37°C with shaking for on-bead-digestion. On following day, 0.5 µg trypsin and 0.05 mAu Lys-C were added to the beads and incubated for 2 hours. Digested peptides in the supernatant were collected into a fresh tube and the beads were 2X washed with HPLC-grade water and pooled with the peptides. Pooled peptides were centrifuged at 16,000 g for 10 minutes and filtered using BioPureSPN columns (Nest Group, Cat. # C100500), pre-wetted with 0.1% trifluoroacetic acid, and spun at 3,000 g for 2 minutes. Filtered peptides were acidified to 2% formic acid, dried using a speed vac and stored at -80°C. Peptides were resuspended in 13 µl of 98/2 buffer A (water + 0.1% formic acid) / buffer B (99.9% acetonitrile + 0.1% formic acid) and 5 µl peptides were injected for mass spectrometric analysis.

Mass spectrometric analysis-chromatographic separation and label-free quantification: Tryptic peptides were separated by reverse phase nano-HPLC using an Ultimate 3000 RSLCnano System (Thermo Fisher Scientific) with a uPAC Trapping column (Thermo Scientific) and a 50 cm uPAC Neo HPLC column (Thermo Scientific). For peptide separation and elution, mobile phase A was 0.1% formic acid (FA) in water and mobile phase B was 0.1% FA in acetonitrile. Peptides were injected onto the trap column at 10 µL/min for 3 minutes using the loading pump. Initially the nanoflow rate was set at 0.75 µL/min and 2% mobile phase B while the peptides were loaded onto the trap column, at 2.8 minutes the solvent composition was changed to 10% mobile phase B. At 5 minutes the flow rate was dropped to 0.300 µL/min at 12% mobile phase B. A two-step gradient was used from 12% to 20% mobile phase B for 41.8 minutes followed by 20% to 40% mobile phase for 15.9 minutes. The flow rate was then increased to 0.750 µL/min for column washing using seesaw gradients and re-equilibration.

Mass spectrometry analysis was performed on an Orbitrap Eclipse (Thermo Fisher Scientific) operated in data-dependent acquisition mode. The MS1 scans were acquired in Orbitrap at 240k resolution, with a 1 x 106 automated gain control (AGC) target, auto max injection time, and a 375-2000 m/z scan range. MS2 targets were filtered for charge states 2-7, with a dynamic exclusion of 60 seconds, and were accumulated using a 0.7 m/z quadrupole isolation window. MS2 scans were performed in the ion trap at a turbo scan rate following higher energy collision dissociation with a 35% normalized collision energy. MS2 scans used a 1 x 104 AGC target and 35 ms max injection time.

### Mass spectrometry data analysis-protein identification and data filtering for sample comparison

Raw MS data files were processed for protein identification and label-free quantification (LFQ) by MaxQuant (version 2.5.1.0) (Tyanova S et al., 2016a) using the Human SwissProt canonical sequence database (UP000005640, downloaded February 2024). The following parameters were used: specific tryptic digestion up to four missed cleavages, variable modification search for upto 5 modification per peptide including carbamidomethyl cysteine, protein N-terminal acetylation, and methionine oxidation, default match between run parameters and label-free quantification with minimum ratio count of 1. Only unmodified, oxidized or N-term acetylated unique peptides were used for protein quantification. The mass spectrometry proteomics data have been deposited to the ProteomeXchange Consortium (http://proteomecentral.proteomexchange.org) via the PRIDE partner repository with the dataset identifier PXD056257.

Quantified protein intensity data from MaxQuant were imported and analyzed by Perseus (version 2.0.11.0) (Tyanova et al., 2016b). First the data were filtered based on categorical column to remove proteins labeled as “Only identified by site”, “Reverse”, and “Potential contaminant”. Then Intensity values were log2 transformed and replicates were grouped in Categorical annotation rows. For this grouping the N-term and C-term miniTurbo-tagged KEAP1 samples were combined into either VVD-065 treated or DMSO treated groups to detect most robust changes consistent across alternatively tagged KEAP1 cell lines. Data were further filtered to remove proteins without quantitative data (valid values) in all eight replicates (four from mT-KEAP1 and four from KEAP1-mT) in at least one group (VVD-065 treated or DMSO treated). Missing values were replaced based on normal distribution with a width of 0.3 and a downshift of 1.8 from the standard deviation for the total matrix. Compound (VVD-065) treated vs control (DMSO) group were compared using two-sided t-test with FDR of 0.05 and S0 of 0 with the default permutation-based FDR correction for multiple t-tests. Volcano plot was generated by GraphPad Prism 10.

